# Benchmarking Machine Learning and Automated Image Analysis for Organelle Quantification

**DOI:** 10.64898/2026.02.12.705524

**Authors:** Chloé Daul, Pierre Tournier, Shukry J. Habib

## Abstract

Quantitative organelle analysis is highly sensitive to image-processing choices, limiting reproducibility across microscopy studies. Here, we systematically compare automated, interactive machine learning, and deep learning–based pipelines for lipid droplet and mitochondrial quantification in live human osteosarcoma cells imaged by fluorescence microscopy and label-free holotomography. Using standardized downstream feature extraction, we evaluated script-based workflows (Fiji, Python), a modular platform (CellProfiler), interactive machine learning (ilastik), and pretrained deep learning models. Lipid droplet segmentation was qualitatively consistent across approaches; however, droplet counts, and size distributions varied substantially between pipelines and imaging modalities, with ilastik reducing background-driven detections and improving cross-modality agreement. In contrast, mitochondrial quantification proved highly sensitive to segmentation and skeletonization choices, particularly in holotomography where global intensity-threshold–based methods failed to capture network structure. Based on these cross-pipeline comparisons, we demonstrate how organelle- and modality-specific benchmarking can guide pipeline selection, illustrated by the analysis of metabolic perturbations affecting lipid droplets and mitochondria. Together, these results highlight modality- and morphology-dependent limitations in common analysis pipelines and provide practical guidance for selecting robust, reproducible strategies for quantitative organelle imaging.

## Introduction

Quantitative image analysis is essential for extracting meaningful biological information from microscopy data. However, reproducible quantification of intracellular structures often relies on complex image processing pipelines involving multiple steps, including preprocessing, segmentation, post-processing, and feature extraction^1^. These steps are frequently sensitive to parameter choices and software implementation, making it challenging to compare results across studies and increasing the risk that biologically meaningful differences are confounded with analysis-dependent artifacts. This highlighs the need for automated and standardized analysis workflows that minimize user-dependent variability and improve reproducibility.

Lipid droplets and mitochondria are central organelles in cellular metabolism and stress responses. Quantitative changes in their abundance, morphology, and network organization are commonly used as indicators of cellular phenotype^2–13^. Lipid droplets are dynamic intracellular organelles that store neutral lipids and help maintain metabolic homeostasis, with changes in droplet size and number reflecting shifts in cellular lipid metabolism and energy balance^14–16^. Meanwhile, mitochondria form highly interconnected and adaptable networks whose morphology is tightly linked to metabolic activity, oxidative stress, and overall cell health^17–20^. Quantifying these structures is therefore of broad interest, but remains technically challenging due to their heterogeneity in size, shape, and spatial organization. Errors in segmentation or network reconstruction can propagate directly into downstream biological interpretation. In particular, mitochondrial network analysis requires not only reliable segmentation but also robust skeletonization and downstream network quantification.

Fluorescence microscopy is widely used for organelle imaging due to its molecular specificity, with probes such as BODIPY for lipid droplets and MitoTracker for mitochondria^13,21–23^. However, fluorescence images can be affected by background signal, out-of-plane fluorescence, non-specific labeling, photobleaching, and variability in excitation intensity, all of which can introduce signal heterogeneity. These effects are particularly pronounced when three-dimensional stacks are reduced using maximum intensity projection, in which only the brightest pixel values along each viewing axis are retained^24^, potentially biasing quantitative measurements toward high-intensity structures. Label-free holotomographic (HT) microscopy provides an alternative by imaging live cells through three-dimensional refractive index maps, avoiding the use of fluorescent dyes and thereby reducing dye-associated toxicity^24–27^. While holotomographic imaging offers intrinsic contrast for certain intracellular structures, the indirect relationship between refractive index and biological identity complicates segmentation, particularly for filamentous organelles such as mitochondria^28^.

A wide range of image analysis software is currently used for quantitative biological microscopy. This includes open-source platforms such as Fiji/ImageJ, CellProfiler, ilastik, and Python-based workflows, as well as commercial or application-specific tools such as TomoAnalysis^29–37^. The open-source platforms are widely adopted in the life sciences because they are accessible, flexible, and suitable for both exploratory and high-throughput analyses. Fiji/ImageJ and CellProfiler have become long-standing standards for classical intensity-based segmentation and object quantification. They offer many community-developed plugins and modular pipelines that support reproducible image analysis across biological applications. In contrast, ilastik provides an interactive machine learning framework that enables supervised pixel classification from user-annotated examples. This reduces the need for manual threshold selection while maintaining interpretability. More recently, deep learning–based approaches have been introduced for automated organelle segmentation. These include pretrained convolutional neural networks such as MitoSegNet and MoDL for mitochondrial analysis^35,36^. By learning complex spatial and intensity patterns directly from large training datasets, these models can capture structures that are difficult to segment with classical thresholding or feature-based methods and can reduce user intervention once trained. However, their performance depends on how closely new images resemble the training data, and adaptation may be required for different imaging conditions or biological systems. Recent work has also demonstrated the potential of deep learning for mitochondrial segmentation in label-free holotomographic microscopy, for example using U-Net–based models to visualize and segment endothelial cell mitochondria^28^. While such approaches highlight the promise of data-driven segmentation for holotomographic imaging, they typically focus on single subcellular structure (e.g., organelles) or specific cell types and do not address how segmentation choices impact downstream quantitative network metrics.

Several studies have benchmarked image analysis pipelines for lipid droplet and mitochondria detection and reported substantial variability in segmentation outcomes across Fiji/ImageJ, CellProfiler, ilastik, and related tools^30,38–40^. These studies show that results are sensitive to threshold selection and preprocessing, and that downstream measurements of object number and size can differ markedly between methods. Similar effects have been observed for mitochondrial segmentation and network analysis, where quantitative metrics depend strongly on both segmentation strategy and downstream network construction, particularly for topology-based features^40^.

Despite this growing body of work, systematic comparisons spanning multiple organelles, imaging modalities, and commonly used analysis platforms remain limited. To our knowledge, no study has evaluated label-free holotomographic microscopy alongside fluorescence imaging while also examining how different segmentation paradigms affect both lipid droplet morphology and mitochondrial network organization within the same experimental framework. To address this gap, we developed standardized and automated image analysis workflows for lipid droplet and mitochondrial quantification and applied them to paired fluorescence and holotomographic live-cell images. We evaluated a representative set of open-source, commercial, and deep learning–based tools (Fiji, Python, CellProfiler, ilastik, TomoAnalysis, MitoSegNet/MitoA, MitoAnalyzer, and MoDL), covering global-intensity-based thresholding, supervised machine learning, and pretrained neural network approaches. Using consistent downstream quantification across pipelines, we systematically assessed how segmentation strategy, imaging modality, and analysis implementation influence quantitative measurements of organelle abundance, morphology, and mitochondrial network organization. Together, this work provides evidence-based guidance for selecting analysis pipelines tailored to organelle morphology and imaging modality and highlights key considerations for improving reproducibility in quantitative cell imaging studies.

## Results & Discussion

### Establishing a unified experimental and image analysis framework

To enable direct comparison between different imaging analysis workflows, we standardized the acquisition and preprocessing steps across modalities (Figure 1). Live TE85 cells were imaged using label-free holotomographic microscopy and stained with BODIPY for lipid droplets and MitoTracker for mitochondria for fluorescence microscopy imaging. Then, three-dimensional image stacks were reconstructed and reduced to two-dimensional representations appropriate for downstream analysis. In total, N = 104 fields of view were acquired, corresponding to n = 1291 imaged cells, with between 1 and 33 cells per image.

**Figure 1.**
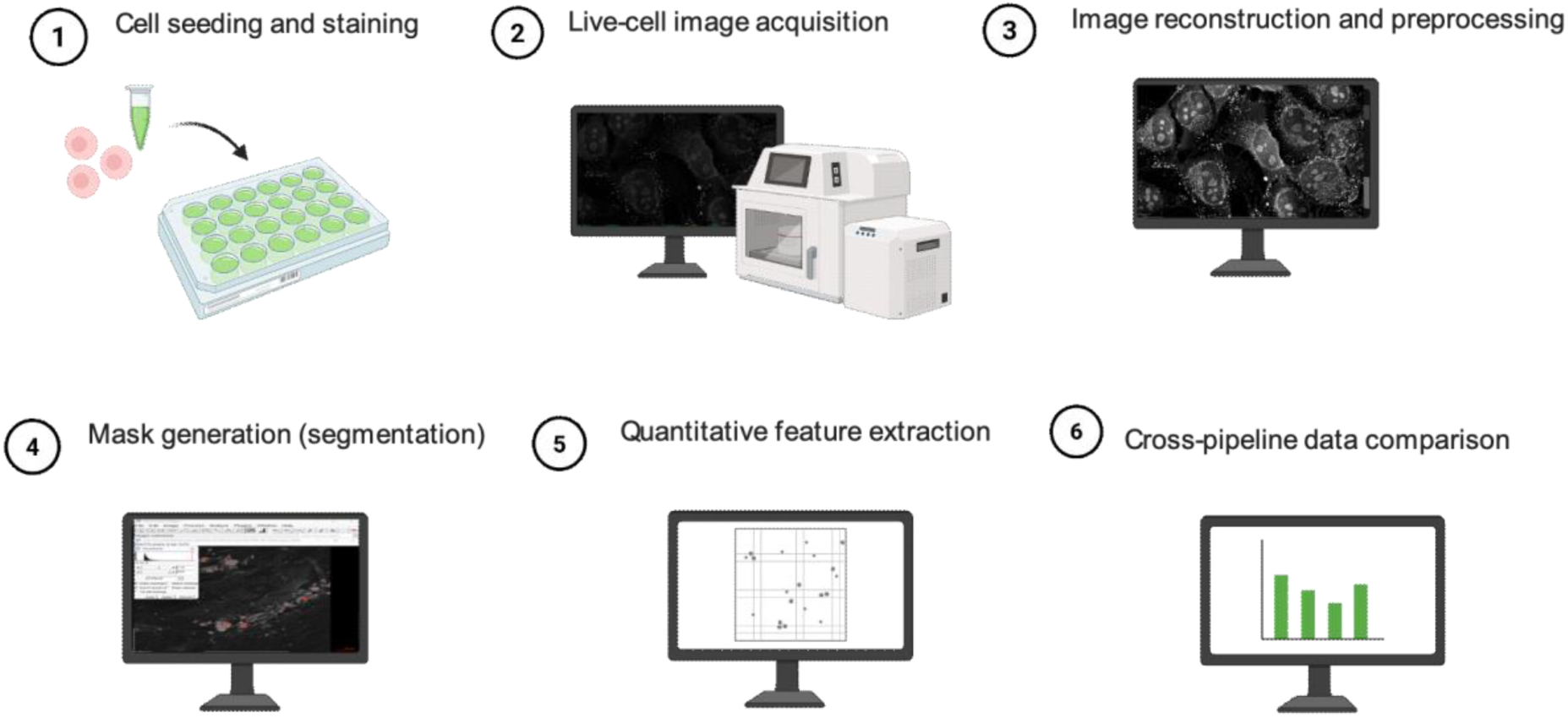
Overview of the experimental and image analysis workflow. Live TE85 cells were imaged using label-free holotomographic microscopy and stained with BODIPY for lipid droplets and MitoTracker for mitochondria for fluorescence microscopy imaging. Image stacks were processed, segmented, and quantified using multiple analysis pipelines, followed by cross-pipeline comparison of quantitative features.

Depending on the software, segmentation and quantitative analysis were either performed within a single environment (end-to-end pipelines such as Fiji, Python, CellProfiler, TomoAnalysis, and MitoAnalyzer) or separated into two steps, with segmentation masks exported for downstream feature extraction (ilastik, MitoSegNet/MitoA, and MoDL). Across all approaches, the general workflow consisted of image preprocessing, segmentation to generate binary masks, identification of individual objects, and extraction of morphological or network-related features (Figure 1). For mitochondria, binary masks were further skeletonized to enable network-based quantification. This approach allowed us to isolate how differences in segmentation strategy and implementation propagate to quantitative measurements.

Within this unified framework, segmentation was performed using distinct strategies depending on the software. Fiji- and Python-based workflows used manually defined global intensity thresholds to generate binary segmentation masks, with thresholds applied with the default SetThreshold function in Fiji and implemented in Python by retaining pixels with intensities above the chosen threshold. CellProfiler used global intensity thresholding implemented via an Otsu method^41^ that automatically determines the threshold from the image intensity distribution. Although MitoAnalyzer also provides automated threshold detection, we instead applied manually defined global threshold to control for parameter selection and ensure that observed differences reflected segmentation strategy rather than tool-specific optimization. In contrast, ilastik used manually annotated training data to classify pixels and directly generate segmentation masks that were exported for downstream analysis, allowing the segmentation to adapt to modality-specific image characteristics. Deep learning–based approaches (MitoSegNet and MoDL) likewise generated segmentation masks directly but relied on pretrained neural networks that learn image features from large external datasets to identify organelle regions. As a result, deep learning segmentation does not depend on user-defined intensity thresholds and instead produces masks based on learned patterns of shape, texture, and intensity, enabling fully automated inference.

For lipid droplets, analysis was performed on two-dimensional maximum-intensity projections derived from three-dimensional fluorescence (BODIPY) and holotomographic stacks (Figure 2). This dimensionality reduction reflects common practice in high-throughput lipid droplet analysis but may amplify high-intensity structures at the expense of weaker signals. For mitochondria, fluorescence (MitoTracker) stacks were similarly reduced to maximum-intensity projections, whereas holotomographic data were represented by a single best-focus slice (Figure 3), as filamentous mitochondrial structures were more reliably resolved in focused refractive-index planes than in projected volumes.

**Figure 2.**
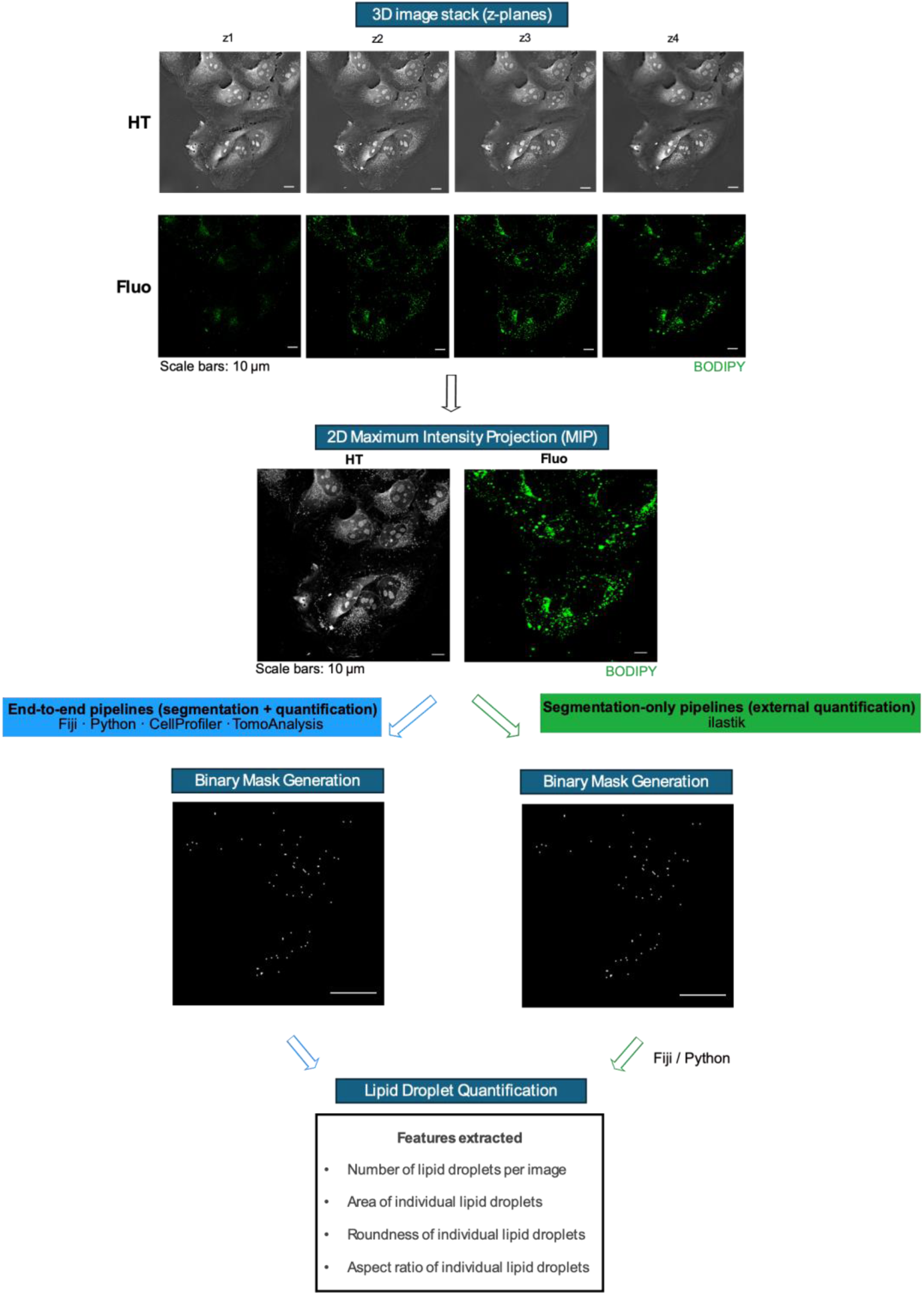
Overview of lipid droplet image processing, segmentation, and quantification pipelines. Representative holotomographic and corresponding fluorescence images acquired using a TomoCube microscope are shown as selected optical sections (z-planes) from three-dimensional image stacks. For clarity, only a subset of z-planes is displayed.

**Figure 3.**
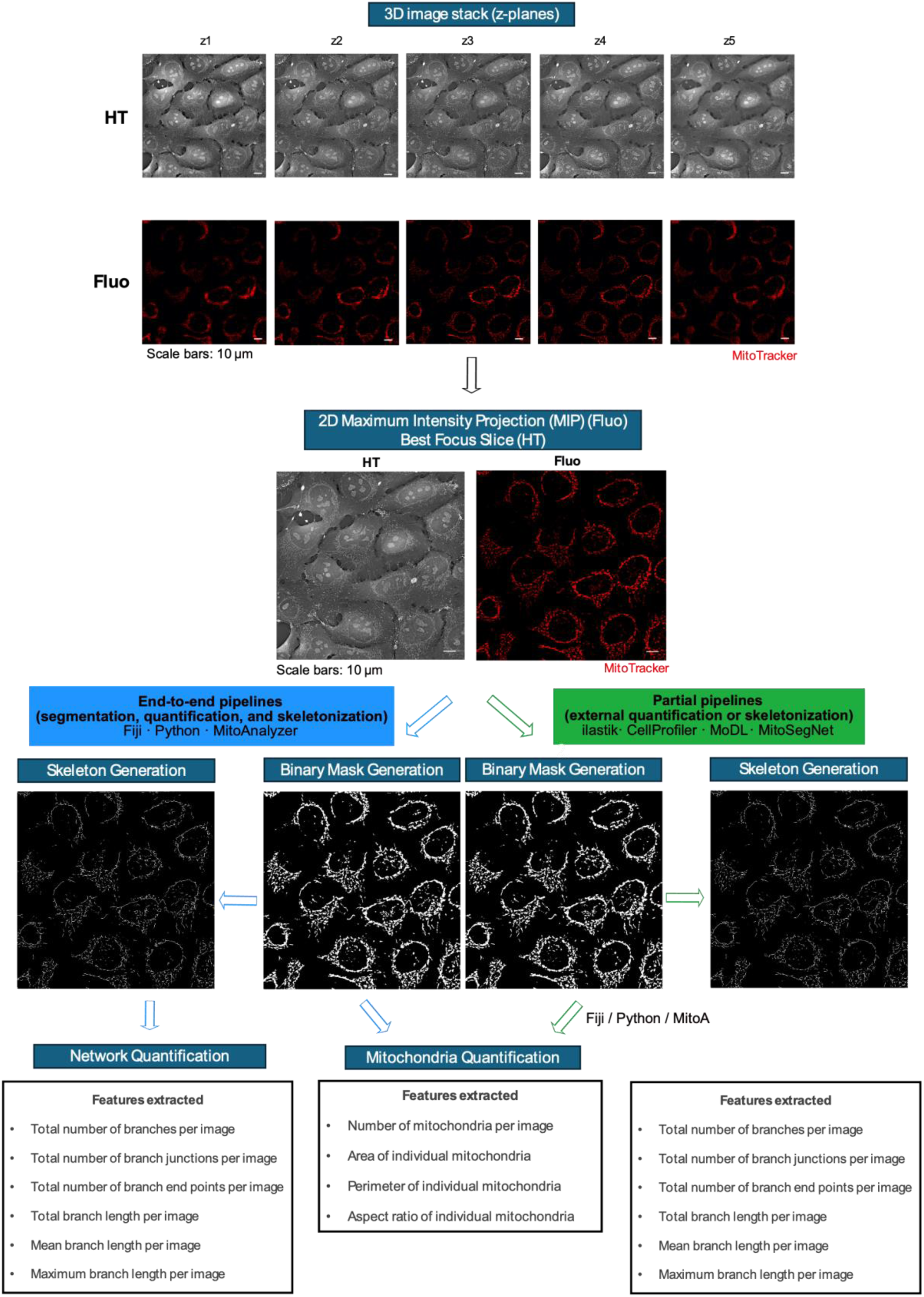
Overview of mitochondrial image processing, segmentation, and network quantification pipelines. Representative holotomographic and corresponding fluorescence images acquired using a TomoCube microscope are shown as selected optical sections (z-planes) from three-dimensional image stacks. For clarity, only a subset of z-planes is displayed.

Across all pipelines, we extracted identical quantitative features, including object counts, area, roundness, and aspect ratio for lipid droplets, and area, perimeter, aspect ratio, and network-related metrics (branch number, junctions, endpoints, and branch lengths) for mitochondria. These features are commonly used to capture biologically relevant differences in lipid droplet abundance and morphology, as well as mitochondrial structure and network organization^20,27,42^.

For ilastik-based segmentation, classifiers were trained on manually annotated subsets of images (10 images per modality for lipid droplets; 10 fluorescence and 54 holotomographic images for mitochondria) and then applied to the full dataset. The larger training set for holotomographic mitochondria reflects the increased heterogeneity and lower intrinsic contrast of these images, requiring additional examples to capture relevant features. Pretrained deep learning models (MitoSegNet and MoDL) were applied directly without retraining or fine-tuning, reflecting their use with default pretrained weights and inference settings as provided by the original developers, without domain adaptation or parameter optimization for our imaging conditions, thereby representing typical plug-and-play deployment by end users. For threshold-based pipelines, segmentation parameters were defined on representative images and subsequently applied uniformly across all corresponding datasets.

Schematics overview of lipid droplets holotomographic (Figure 2; Supplementary Figure S1a-c) and fluorescence workflows (Figure 2; Supplementary Figure S2a-d) and mitochondria holotomographic (Figure 3; Supplementary Figure S3a-c) and fluorescence workflows (Figure 3; Supplementary Figure S4a-d) are provided.

### Lipid droplet detection and quantitative analysis across pipelines

We first compared lipid droplet appearance across imaging modalities by visual inspection of representative images and corresponding segmentation masks (Figure 4a-b). Fluorescence images stained with BODIPY displayed a higher number of lipid-associated structures than holotomographic images (Figure 4b-c), indicating modality-dependent detection. Some fluorescent structures lacked counterparts in holotomographic images (Figure 4a-b, red arrows). However, a subset of lipid droplets was consistently detected across both modalities (Figure 4a-b, blue arrows), supporting the ability of holotomographic imaging to capture lipid-rich regions through intrinsic refractive index contrast^43^. Further separation of modality-specific detections would require full three-dimensional analysis and was not pursued here.

**Figure 4.**
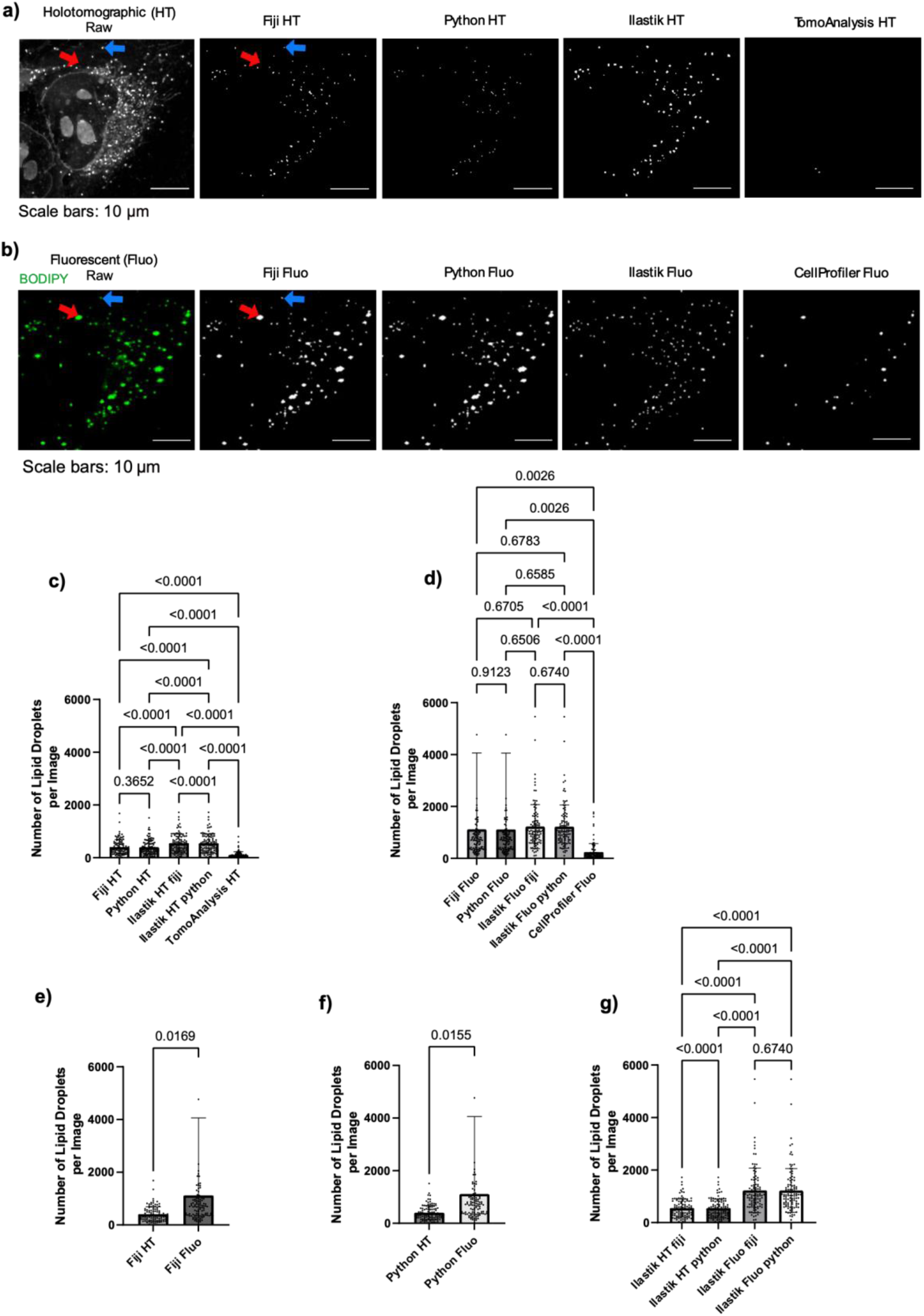
Comparison of lipid droplet segmentation and quantitative analysis across image analysis pipelines. (a) Maximum intensity projections (MIP) of representative holotomographic (HT) raw images used as input for segmentation, together with the corresponding binary masks generated using Fiji, Python, and ilastik. (b) Maximum intensity projections (MIP) of representative fluorescence (Fluo, BODIPY) raw images used as input for segmentation, together with the corresponding binary masks generated using Fiji, Python, ilastik, and CellProfiler. Red arrows highlight a structure detected as a lipid droplet in fluorescence but not clearly visible in the holotomographic image, illustrating modality-dependent detection, while blue arrows indicate a lipid droplet consistently detected across imaging modalities and segmentation methods. (c) Number of lipid droplets per image quantified from holotomographic (HT) images using different analysis pipelines (Fiji, Python, ilastik, and TomoAnalysis). (d) Number of lipid droplets per image quantified from fluorescence (Fluo, BODIPY) images using different analysis pipelines (Fiji, Python, ilastik, and CellProfiler). (e) Paired comparison of lipid droplet counts per image obtained from holotomographic and fluorescence images using the Fiji-based pipeline. (f) Paired comparison of lipid droplet counts per image obtained from holotomographic and fluorescence images using the Python-based pipeline. (g) Comparison of lipid droplet counts per image obtained using ilastik-based segmentation across imaging modalities (HT and Fluo) and downstream quantification environments. Statistical comparisons between pipelines or conditions are indicated above each plot. *N*=104 images, each image contains 1-33 cells. Statistical analyses: One-way ANOVA.

Segmentation outcomes differed across analysis pipelines. In holotomographic images, TomoAnalysis identified fewer droplets than the other methods, likely reflecting conservative default parameterization (Figure 4a). In fluorescence images, Fiji- and Python-based workflows using manually defined global intensity thresholds closely followed the raw signal and detected many low-intensity structures (Figure 4b). CellProfiler-derived masks preferentially detected brighter and larger droplets and showed reduced sensitivity to smaller or weaker signals (Figure 4b). Ilastik-based supervised pixel classification suppressed low-intensity or ambiguous detections and produced segmentation masks that more closely resembled those obtained from holotomographic images when applied to fluorescence data (Figure 4b).

Quantitative comparisons revealed substantial variability in the number of lipid droplets detected per image across pipelines and imaging modalities (Figure 4c-d). Fluorescence-based analyses yielded higher droplet counts than holotomographic analyses across pipelines (Figure 4e-f). TomoAnalysis and CellProfiler detected fewer droplets than Fiji- and Python-based workflows, while the latter two produced similar droplet counts reflecting their shared thresholding strategy (Figure 4c-d). Using ilastik-derived segmentation masks, lipid droplet counts were similar whether downstream quantification was performed in Fiji or in Python for fluorescence and holotomographic images (Figure 4g).

Morphological analysis further revealed pipeline-dependent effects. Mean lipid droplet area differed significantly between imaging modalities within the same software (Supplementary Figure S5a-e), whereas shape-related features such as roundness and aspect ratio (a measure of object elongation calculated as 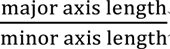) were broadly comparable across pipelines (Supplementary Figure S5f-o). Within-image variability in lipid droplet area was generally higher for fluorescence-based analyses than for holotomographic analyses, except for ilastik-based segmentation, which showed similar variability across modalities (Supplementary Figure S6a-o). Consistent with visual inspection, fluorescence images exhibited a broader range of lipid droplet sizes (Supplementary Figure S5a-b).

These pipeline-dependent differences have direct biological implications. In fluorescence images, segmentation strategy strongly influences lipid droplet abundance estimates. More permissive global thresholding approaches used in Fiji and Python increase detection sensitivity but are more susceptible to background signal and low-intensity detections, whereas more conservative approaches such as CellProfiler preferentially detect brighter and larger droplets and may underestimate lipid droplet abundance. In holotomographic images, conservative detection strategies, particularly those implemented in TomoAnalysis, yielded lower droplet counts, reflecting reduced sensitivity to smaller or lower-contrast lipid droplets. Because metabolic^44^ or phenotypic perturbations^45^ can alter lipid droplet size or intensity without necessarily changing droplet number, such pipelines may misinterpret morphological remodeling as lipid droplet loss.

Overall, these results demonstrate that lipid droplets tolerate a broad range of segmentation strategies due to their simple morphology and stable shape features. Global intensity-threshold workflows are sufficient to detect strong lipid droplet phenotypes. Learning-based segmentation improves cross-modality agreement when suppression of low-intensity or ambiguous structures is prioritized.

### Mitochondria detection and quantitative analysis across pipelines

We next extended our comparative analysis to mitochondria, a structurally complex organelle whose filamentous network organization poses additional challenges for segmentation and quantification. Visual inspection revealed substantially larger discrepancies between mitochondrial segmentation pipelines than those observed for lipid droplets, particularly in holotomographic images (Figure 5A–B vs. Figure 4A–B). Comparison between imaging modalities further highlighted remaining limitations. Some mitochondria visible in fluorescence images were only partially captured in holotomographic-based segmentations (Figure 5A–B, blue arrow), whereas other structures detected as mitochondria in holotomographic masks were absent from fluorescence images and likely correspond to non-mitochondrial features such as cell boundaries or thin membrane protrusions (Figure 5a-b, red arrow).

**Figure 5.**
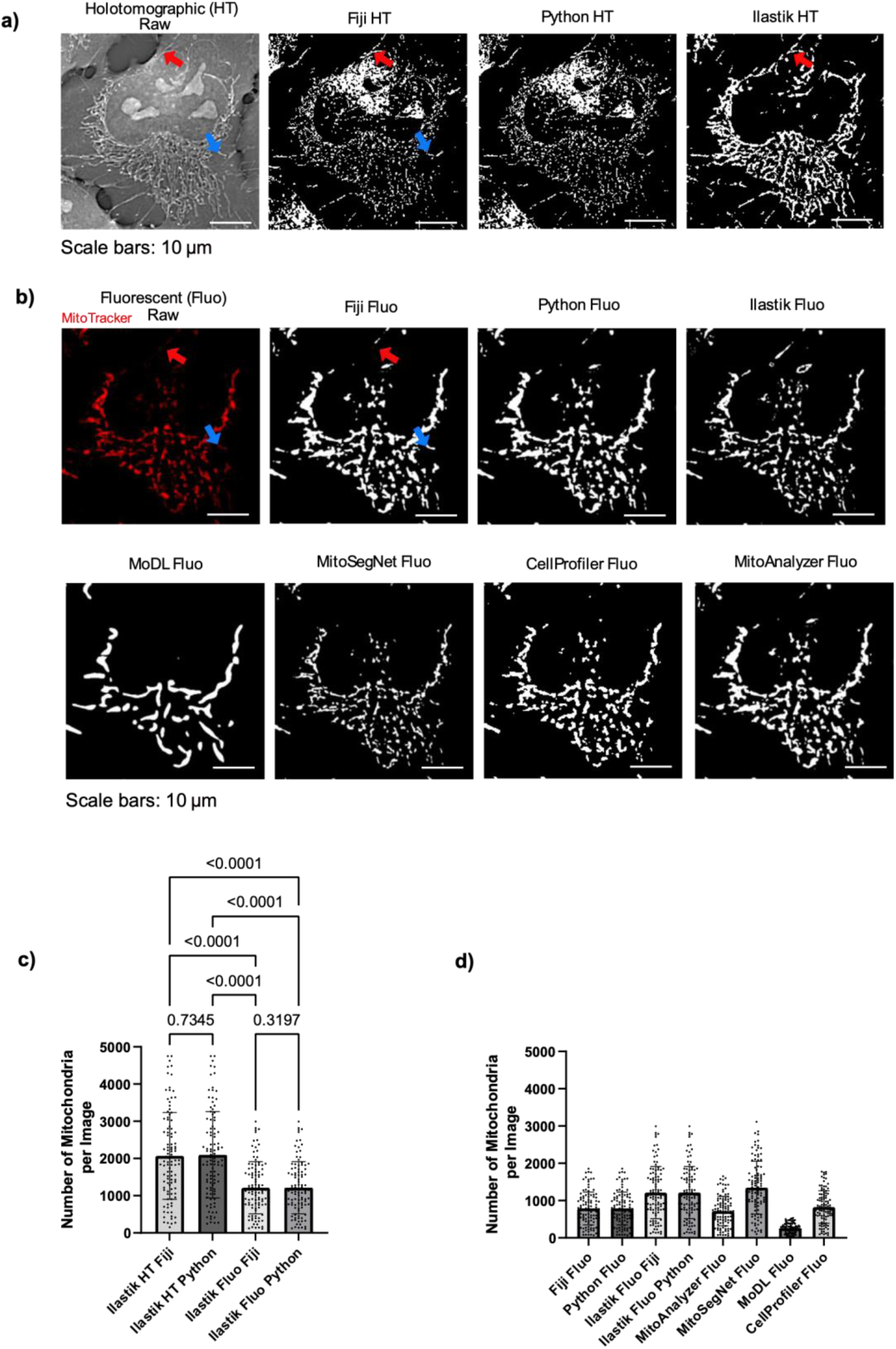
Comparison of mitochondrial segmentation and quantitative analysis across image analysis pipelines. (a) Best-focus holotomographic (HT) images used as input for segmentation, together with the corresponding binary masks generated using Fiji, Python, and ilastik. (b) Maximum intensity projections (MIP) of corresponding fluorescence (Fluo, MitoTracker) images used as input for segmentation, together with the corresponding binary masks generated using Fiji, Python, ilastik, MoDL, MitoSegNet, CellProfiler, and MitoAnalyzer. Red arrows highlight a structure detected as mitochondria in holotomographic-based segmentation but not visible in the fluorescence image, likely corresponding to a non-mitochondrial cellular structure (e.g., cell boundary), illustrating a false-positive detection in holotomographic-based analysis, while blue arrows indicate a mitochondrial filament consistently detected across imaging modalities and segmentation methods. (c) Comparison of the number of mitochondria per image obtained using ilastik-based segmentation across imaging modalities (holotomographic, HT, and fluorescence, Fluo) and downstream quantification environments. (d) Comparison of the number of mitochondria per image quantified from fluorescence images using different analysis pipelines (Fiji, Python, ilastik, MitoAnalyzer, MitoSegNet, MoDL, and CellProfiler). Masks derived from holotomographic segmentations using threshold-based approaches were excluded from quantitative analysis due to insufficient segmentation quality. Only p-values > 0.01 are shown in the graph; complete statistical comparisons are reported in in Supplementary Table 1. Differences between conditions were assessed using repeated-measures one-way ANOVA (RM one-way ANOVA). *N*=104 images, each image contains 1-33 cells.

For holotomographic images, global intensity threshold–based segmentation implemented in Fiji and Python performed poorly, detecting a heterogeneous mixture of intracellular structures rather than selectively isolating mitochondria (Figure 5a). These biologically implausible masks were unsuitable for downstream network analysis and were therefore excluded from further quantitative comparisons. In contrast, ilastik-based segmentation substantially improved mitochondrial delineation in holotomographic images. Although some non-mitochondrial structures were still detected (Figure 5a, red arrow (ilastik HT)), the resulting masks more closely resembled those obtained from fluorescence images, indicating that learning-based approaches are better suited to handling the lower contrast and structural complexity of mitochondria in label-free holotomographic data.

For fluorescence images, several pipelines produced visually similar segmentation masks (Figure 5b). Fiji-, Python-, and MitoAnalyzer-based approaches generated identical masks because they relied on the same intensity-based segmentation parameters. Other fluorescence-based pipelines showed greater variability: MoDL frequently under-segmented mitochondrial networks, whereas ilastik and MitoSegNet better preserved mitochondrial continuity and more closely matched the raw fluorescence signal (Figure 5b). CellProfiler yielded reasonable segmentations (Figure 5b).

Quantitative analysis revealed substantial variation in mitochondrial counts across pipelines (Figure 5c-dD). Under-segmentation reduced object counts by merging structures (Figure 5d; MoDL), while differences in downstream processing led to divergent counts even when identical masks were used. In our analysis, Fiji and Python quantified mitochondrial number by connected-component labeling of the binary masks without applying a minimum size threshold (size = 0 to ∞). In contrast, MitoAnalyzer applied a minimum area threshold (≥0.06 µm²), reducing both mitochondrial counts and total area despite identical initial segmentation (Figure 5d).

As observed for lipid droplets (Figure 4g), ilastik-based segmentation yielded comparable mitochondrial counts whether downstream quantification was performed using Fiji or Python, for both fluorescence and holotomographic images (Figure 5c-d), indicating robustness to the choice of downstream analysis environment.

Analysis of mitochondrial morphology further revealed pipeline-dependent effects, particularly for boundary-sensitive metrics such as mean area, perimeter, and aspect ratio (Supplementary Figure S7a-f) Variability metrics, assessed as the per-image standard deviation (Supplementary Figure S7a-f), were influenced more strongly by the analysis pipeline than by imaging modality alone, indicating that some workflows amplify apparent heterogeneity while others suppress it. This has important implications for studies of mitochondrial dynamics or stress responses, where changes in variability are often biologically interpreted^46,47^.

Together, these results demonstrate that mitochondrial detection and object-level quantification are highly dependent on the chosen analysis pipeline. Global intensity threshold–based approaches, such as those implemented in Fiji and Python, offer speed, transparency, and reproducibility, but were insufficient for reliable mitochondrial segmentation in holotomographic images and remained sensitive to noise and intensity variations in fluorescence data. Learning-based methods, including ilastik and deep learning–based approaches, better preserved mitochondrial continuity and complex morphology, particularly in label-free holotomographic microscopy. These improvements come with practical trade-offs, including manual training requirements for ilastik and increased computational costs. Deep learning–based approaches represent a promising direction for label-free mitochondrial segmentation^28^, but their performance and generalizability remain strongly dependent on training data and experimental context, consistent with the underperformance of MoDL in our dataset.

### Mitochondrial network skeletonization and quantitative analysis

Mitochondrial function is closely linked to the organization and connectivity of the mitochondrial network. Thus, we next examined how differences in segmentation pipelines propagate to downstream skeletonization and network-based quantification. Skeletonization was performed on binary mitochondrial masks and therefore largely reflected trends already observed at the segmentation stage (Figure 6a-b). Pipelines that produced more extensive or continuous mitochondrial masks yielded denser skeleton networks, whereas pipelines that under-segmented mitochondria resulted in fewer or more fragmented skeletons, indicating that segmentation quality directly influence downstream network representations.

**Figure 6.**
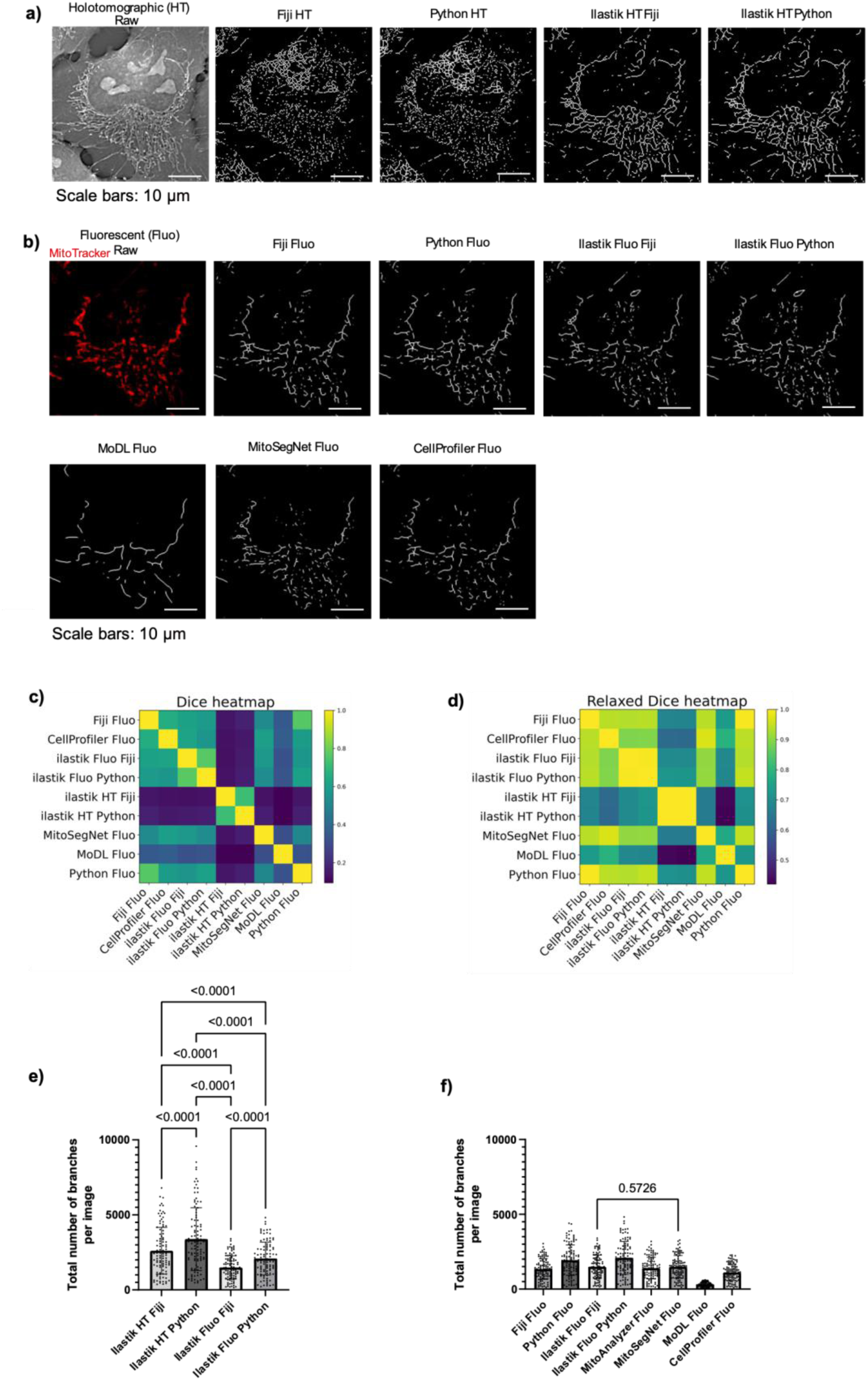
Comparison of mitochondrial network skeletonization and quantitative analysis across image analysis pipelines. (a) Representative mitochondrial skeletons derived from holotomographic (HT) images following segmentation using different analysis pipelines (reproduced from Figure 5 for consistency). (b) Representative mitochondrial skeletons derived from fluorescence (Fluo, MitoTracker) images following segmentation using different analysis pipelines (reproduced from Figure 5 for consistency). (c) Heatmap of pairwise Dice coefficients computed between mitochondrial skeletons generated by different analysis pipelines. The Dice coefficient quantifies the exact pixel-wise overlap between two skeletons, with values ranging from 0 (no overlap) to 1 (perfect overlap). (d) Heatmap of relaxed Dice coefficients computed using a tolerance of ±2 pixels. In this relaxed metric, skeletons were symmetrically dilated prior to overlap calculation to account for small spatial offsets between pipelines. Differences between conditions were assessed using repeated-measures one-way ANOVA (RM one-way ANOVA) (e) Comparison of the total number of mitochondrial network branches per image obtained using ilastik-based segmentation across imaging modalities (holotomographic, HT, and fluorescence, Fluo) and downstream skeletonization environments. (f) Comparison of the total number of mitochondrial network branches per image quantified from fluorescence images using different analysis pipelines (Fiji, Python, ilastik, MitoAnalyzer, MitoSegNet, MoDL, and CellProfiler). Skeletons derived from holotomographic segmentations using threshold-based approaches were excluded from quantitative analysis due to insufficient segmentation quality for reliable network quantification. Selected statistical comparisons are indicated directly on the plots, while the complete set of statistical comparisons is reported in Supplementary Table 8. *N*=104 images, each image contains 1-33 cells.

For fluorescence images, visual inspection revealed broadly similar mitochondrial network patterns across most pipelines (Figure 6b). Continuous filamentous structures were preserved in skeletons derived from Fiji, Python, ilastik, MitoSegNet, CellProfiler, and MitoAnalyzer segmentations. In contrast, MoDL-derived skeletons appeared sparser and more fragmented, consistent with the reduced mitochondrial coverage observed at the mask level (Figure 6b vs Figure 5b). This underlines that incomplete segmentation leads to fragmented skeletons rather than reflecting true differences in mitochondrial network complexity.

Skeletons derived from holotomographic images highlighted additional challenges. Here, skeletons generated using global intensity-threshold–based segmentation approaches (Fiji and Python) were noisy and biologically implausible and were therefore included for qualitative comparison only (Figure 6a). These skeletons were excluded from quantitative network analysis. In contrast, ilastik-based holotomographic segmentation produced interpretable skeletons that were suitable for quantitative analysis (Figure 6c), enabling direct comparison of network metrics across pipelines.

To quantitatively assess agreement between mitochondrial skeletons generated by different pipelines, pairwise similarity was evaluated using the Dice coefficient, which measures exact pixel-wise skeleton overlap, and a relaxed Dice coefficient, which allows for small spatial deviations between skeletons (Figure 6c-d; Supplementary Figure S10). Consistent with visual inspection, skeletons derived from holotomographic images generally showed lower similarity to fluorescence-based skeletons, reflecting increased segmentation uncertainty in label-free data. In line with qualitative observations, MoDL-derived skeletons also exhibited reduced similarity to other pipelines, consistent with their sparser and more fragmented network representations. In contrast, most fluorescence-based pipelines showed high mutual agreement, indicating broadly similar mitochondrial network reconstructions. Across all comparisons, relaxed Dice coefficients were systematically higher than standard Dice values (Figure 6c-d), demonstrating that a substantial fraction of disagreement arises from small spatial offsets between skeletons rather than major differences in overall network topology.

Quantitative network analysis revealed substantial variability in the total number of mitochondrial branches detected per image across pipelines (Figure 6e-f). MoDL produced fewer branches than other methods, in line with its tendency to under-segment mitochondrial structures (Figure 6f). In this case, reduced branch counts reflect decreased network continuity at the segmentation stage rather than a biological reduction in mitochondrial branching.

Systematic differences were also observed between skeletonization implementations. Fiji- and Python-based pipelines yielded different branch counts despite being applied to identical segmentation masks (Figure 6f). Python-based skeletonization detected more branches than Fiji-based skeletonization across segmentation methods, including ilastik-derived masks (Figure 6e-f). This demonstrates that skeletonization algorithms alone, independent of segmentation quality, can substantially influence derived network metrics.

Analysis of additional skeleton-based features including the number of branch junctions, number of branch endpoints, total branch length, mean branch length, and maximum branch length generally mirrored the trends observed for branch counts (Supplementary Figure S9a-j). Pipelines producing denser skeletons showed increased numbers of junctions and endpoints and greater total branch length per image, whereas pipelines with sparse or fragmented skeletons yielded reduced values across these metrics (Supplementary Figure S9a-f).

In contrast, mean and maximum branch length varied less across pipelines than branch counts or junction numbers (Supplementary Figure S9g-j). This suggests that while segmentation and skeletonization strongly influence network connectivity and fragmentation, the length of individual mitochondrial segments is comparatively robust. Different network metrics therefore capture distinct aspects of mitochondrial organization and differ in their sensitivity to analysis pipeline choice.

Overall, while accurate segmentation is a prerequisite for generating biologically meaningful skeletons, differences between skeletonization algorithms alone were sufficient to produce divergent network metrics, even when applied to similar or identical masks. Threshold-based segmentation was inadequate for holotomographic images, whereas learning-based methods yielded interpretable skeletons across modalities. Systematic differences between Fiji- and Python-based skeletonization further indicate that network metrics are highly sensitive to downstream analytical choices.

Together, these observations underscore the need to interpret mitochondrial network metrics in conjunction with visual inspection and to standardize both segmentation and skeletonization steps wherever possible. Network-level conclusions should therefore be drawn cautiously, with explicit consideration of how analytical implementation may influence measurements of connectivity, branching, and fragmentation.

To facilitate practical adoption of these findings, we summarize the key characteristics, strengths, and limitations of each evaluated pipeline in Table 1. This overview highlights trade-offs between automation level, user input, generalization to new data, and segmentation robustness, providing a concise reference for selecting appropriate analysis strategies based on experimental goals and technical constraints.

**Table 1.** Comparison of evaluated image analysis pipelines.

| Feature | Fiji / ImageJ | Python (custom) | CellProfiler | ilastik | TomoAnalysis | MitoSegNet | MoDL | MitoAnalyzer |
| --- | --- | --- | --- | --- | --- | --- | --- | --- |
| Segmentation approach | Manual global intensity thresholding | Manual global intensity thresholding | Modular intensity-based pipeline | Supervised pixel classification (ML) | Refractive index-based thresholding (HT) | Pretrained deep learning | Pretrained deep learning | Network analysis (external masks used here) |
| Lipid droplets segmentation (binary mask generation) | ✓ | ✓ | ✓ | ✓ | ✓ | ✗ | ✗ | ✗ |
| Mitochondria segmentation (binary mask generation) | ✓ | ✓ | ✓ | ✓ | ✓ | ✓ | ✓ | ✓ (external masks used here) |
| Mitochondrial network analysis | ✓ | ✓ | ✗ | ✗ | ✗ | ✗ | ✗ | ✓ |
| Internal quantification | ✓ | ✓ | ✓ | ✗ | ✓ (limited) | ✗ (requires MitoA) | ✗ | ✓ |
| Level of automation | Semi-automated | Semi-automated | Automated after setup | Semi-automated | Automated | Fully automated | Fully automated | Automated after setup |
| User input | Thresholds, parameters | Thresholds, scripting | Pipeline design | Manual annotation | Minimal | None (unless retraining) | None (unless retraining) | Binary masks (provided externally in this study) |
| Computational runtime (relative) | Fast | Fast | Moderate | Moderate | Moderate | Slow–moderate | Slow–moderate | Moderate |
| Generalization to new data | High (rule-based) | High (rule-based) | High (rule-based) | Moderate (training-dependent) | Moderate | Limited (requires retraining) | Limited (requires retraining) | High (rule-based) |
| Key strengths | Flexible, visual validation | Customizable, scalable | High-throughput pipelines | Adapts to modality | Integrated HT | Preserves continuity | End-to-end DL | Dedicated network metrics |
| Main limitations | Threshold sensitivity | Coding required | Requires parameter optimization | Requires user training | Limited output metrics | Dataset dependence | High computational cost | Relies on upstream segmentation in this study |

✓ = Yes ✗ = No

### Functional validation of organelle-specific analysis pipelines

To validate our organelle-specific pipeline selection, we applied two well-established perturbations that induce characteristic changes in lipid droplets and mitochondrial organization. Oleic acid treatment was used to promote lipid droplet accumulation^48–50^, while serum starvation was applied to induce mitochondrial network remodeling via metabolic stress ^42,51–53^. These experiments were designed to assess whether the analysis workflows selected based on organelle morphology can detect condition-dependent structural changes.

For lipid droplets, oleic acid treatment served as a robust positive control (Figure 7a-b). Because lipid droplets appear as high-contrast, approximately spherical objects, segmentation was performed using the same Python-based global intensity-thresholding pipeline and identical parameters as in the previous benchmarking analysis, applied uniformly across all images (Oleic acid and Untreated). This lightweight workflow enabled rapid batch processing without additional parameter tuning.

**Figure 7.**
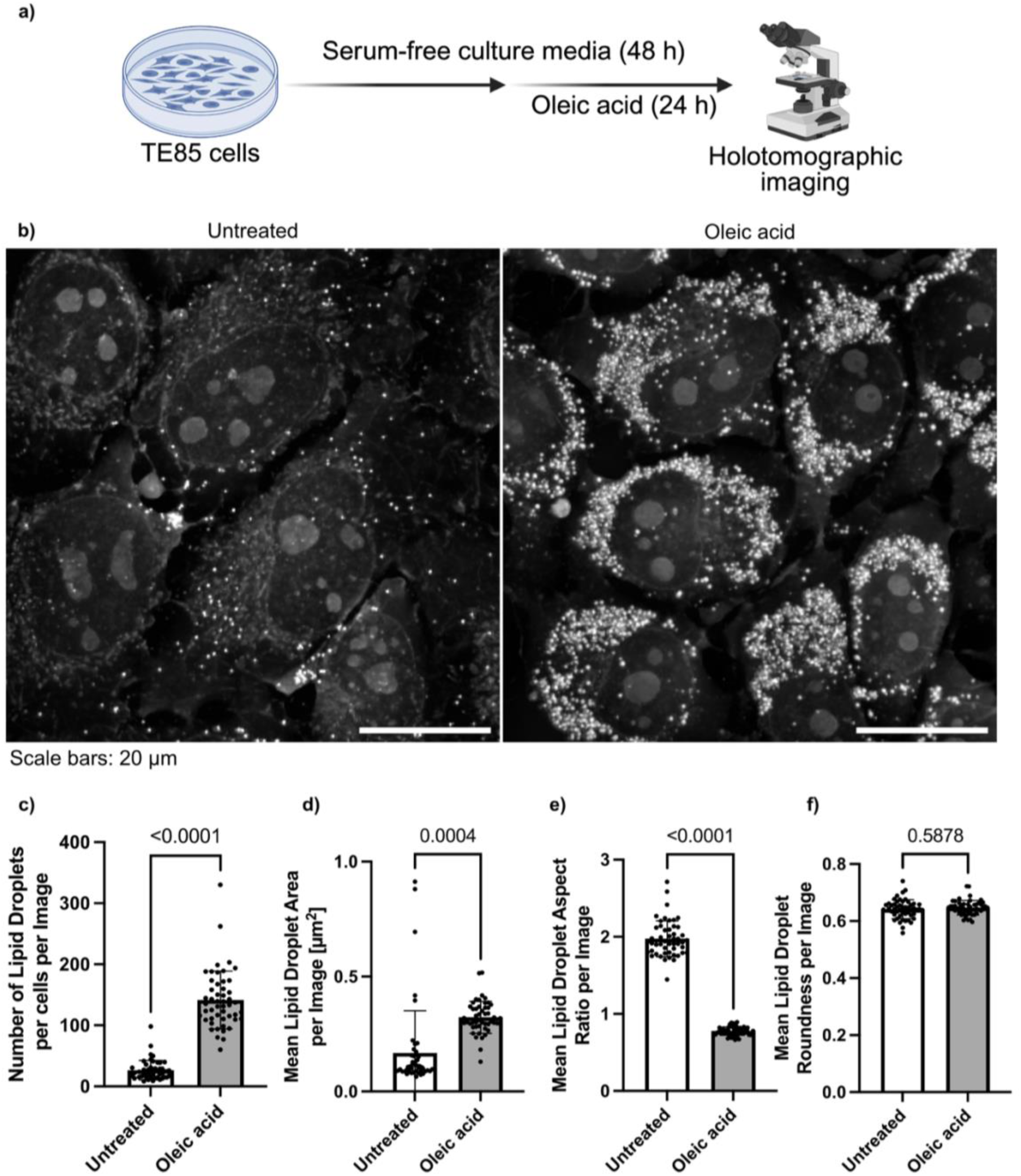
Oleic acid–induced lipid droplet accumulation quantified using a global intensity-threshold–based pipeline. (a) Schematic representation of the experimental workflow. (b) Representative holotomographic images of TE85 cells. (c-f) Quantitative analysis of lipid droplet abundance and morphology following oleic acid treatment compared to untreated cells as control. Lipid droplets were segmented using a Python-based global intensity-thresholding pipeline with identical parameters applied across conditions. (c) Number of lipid droplets per image normalized to the number of cells. (d) Mean lipid droplet area per image. (e) Mean lipid droplet aspect ratio per image. (f) Mean lipid droplet roundness per image. Statistical comparisons between conditions are indicated above each plot. N = 50 images per condition; each image contains 1-29 cells. Statistical analyses: Welch’s t-tests.

Quantitative analysis revealed a marked increase in lipid droplet number per image following oleic acid treatment (Figure 7c), together with significant changes in droplet morphology, including reduced mean droplet area (Figure 7d) and increased aspect ratio (Figure 7e), while roundness remained largely unchanged (Figure 7f). These results demonstrate that this minimal, intensity-based pipeline is sufficient to capture strong lipid droplet phenotypes induced by metabolic perturbation. This finding is consistent with our earlier cross-pipeline comparisons, which showed that lipid droplets can be robustly detected using simple threshold-based approaches due to their relatively uniform shape and strong contrast.

Based on our earlier cross-pipeline comparisons, ilastik better preserved mitochondrial continuity than global intensity-threshold–based methods, particularly in fragmented or low-intensity network regions. For this experiment, we then selected a previously trained ilastik classifier and applied it in batch mode to the new dataset. Binary masks were subsequently analyzed using Python-based morphometric and network quantification. For the direct comparison between a simple threshold-based approach and learning-based segmentation, we analyzed the same dataset using the global intensity-threshold–based Python pipeline.

Using these two pipelines, we compared the mitochondrial network of control (unstarved) and serum-starved (24 h) cells (Figure 8a-b). Quantitative analysis revealed different outcomes depending on the pipeline used. The total number of mitochondria per cell differed between the two pipelines (Figure 8c), with ilastik being able to detect less mitochondria in the starved cells compared to the unstarved ones. This difference was not observed using the Python-based threshold, highlighting the importance of the choice of the analysis pipeline for detecting mitochondrial features. A similar conclusion was made on the mitochondrial area (Figure 8d) and the number of branch endpoints (Figure 8e), where ilastik was able to detect statistical differences between the two groups, unlike Python. Nonetheless, other features showed consistent trends across both pipelines, such as the mitochondrial aspect ratio (Supplementary Figure 11a), the mitochondrial perimeter (Supplementary Figure 11b), the number of branches (Supplementary Figure 11c), the number of branch junctions (Supplementary Figure 11d), the total, mean, and maximum branch length (Supplementary Figure 11e-g). This highlights that pipeline choice strongly influences the detection of mitochondrial remodeling.

**Figure 8.**
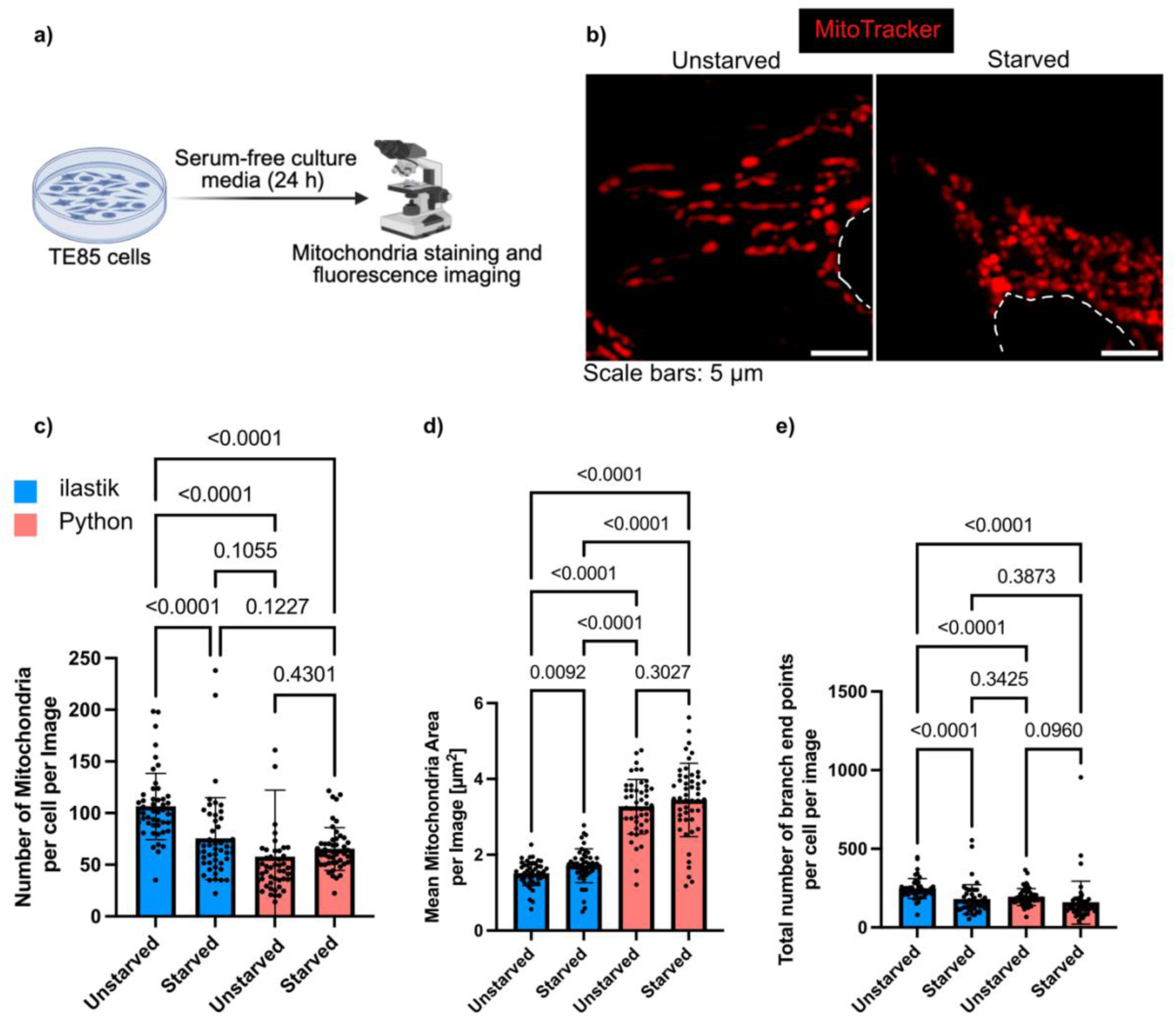
Analysis method choice influences data outcomes for analysis of the mitochondrial network. (a) Schematic representation of the experimental workflow. (b) Representative fluorescent images of mitochondria stained by MitoTracker (red). The white dashed line indicates the edge of the nuclei. (c-e) Quantitative comparison of the number of mitochondria (c), the mean mitochondria area (d), and the number of branches endpoint (e) using ilastik (blue) and Python. N = 50 images per condition; each image contains 1-13 cells. Statistical analyses: Welch’s ANOVA.

Mitochondrial responses to starvation are known to be highly context-dependent, varying with cell type, deprivation regime, and duration, with reports describing both elongation and fragmentation^50–53^. Here, our objective was not to define a universal starvation phenotype, but rather to assess whether the selected analysis pipelines are capable of detecting mitochondrial remodeling between conditions. The internally consistent changes observed across object-level morphology and network topology metrics demonstrate that learning-based segmentation enables sensitive detection of condition-dependent mitochondrial organization in this setting, whereas the Python-based thresholding pipeline shows limitations for certain mitochondrial features.

In summary, these perturbation experiments reinforce our conclusions that organelle morphology should guide pipeline selection. Simple global thresholding is sufficient to capture strong lipid droplet phenotypes, whereas mitochondria require more precise, learning-based segmentation to reveal both morphological and network-level changes. This practical validation illustrates how different analysis strategies lead to distinct detection capabilities and further motivates the use of organelle-specific pipelines for reproducible quantitative imaging.

## Conclusion

Quantitative analysis of lipid droplets and mitochondria is central to many areas of cell biology, as changes in their abundance, morphology, and organization provide key readouts of cellular metabolic state, stress responses, and disease-associated phenotypes^2–13^. Alterations in lipid droplet size and number reflect shifts in lipid storage and energy balance, while mitochondrial morphology and network connectivity are closely linked to bioenergetic function, oxidative stress, and cell viability^14–20^. Robust and reproducible quantification is therefore particularly important in experimental contexts such as drug screening, genetic perturbation, or disease modeling, where subtle changes in organelle structure may be used to infer biological effects.

In this study, we performed a systematic comparison of automated image analysis pipelines for lipid droplets and mitochondrial quantification across fluorescence and label-free holotomographic microscopy. By applying standardized downstream analysis to outputs generated by multiple software tools, we demonstrate that quantitative measurements are strongly influenced by both imaging modality and analysis workflow, even when applied to the same images.

For lipid droplets, most pipelines captured broadly similar qualitative patterns, but quantitative metrics such as droplet number, size, and variability differed substantially across methods. Learning-based segmentation, particularly ilastik, reduced low-intensity or potentially background-related detections and produced results that were more consistent across modalities, albeit at the cost of manual training and increased setup time. Differences in computational performance across lipid droplet pipelines were relatively modest, making workflow selection primarily dependent on the desired balance between sensitivity and robustness rather than execution speed alone.

Mitochondrial analysis proved even more sensitive to pipeline choice. Global intensity-threshold–based segmentation was insufficient for reliable mitochondrial detection in holotomographic images and remained sensitive to noise and intensity variations in fluorescence data. Learning-based approaches better preserved mitochondrial continuity across modalities, enabling downstream morphological and network analysis. Deep learning–based methods such as MitoSegNet and MoDL offered fully automated segmentation but required longer execution times, while ilastik required user-guided training, which was relatively fast for fluorescence images but substantially more time-consuming for holotomographic data due to increased structural complexity.

Importantly, substantial differences in quantitative outcomes arose not only from segmentation quality but also from downstream processing steps, including object identification and skeletonization. In particular, mitochondrial network metrics were significantly affected by the specific skeletonization implementation, demonstrating that variability can be introduced well beyond the initial segmentation stage.

Together, these findings show that no single analysis pipeline performs optimally across both organelle types and imaging modalities. This conclusion was further supported by metabolic perturbation experiments: oleic acid–induced lipid droplet accumulation was readily detected using simple global thresholding, whereas serum starvation–induced mitochondrial remodeling could not be reliably captured with this approach and therefore required learning-based segmentation. These results illustrate how organelle morphology and imaging context inform practical pipeline choice, and how different analysis strategies offer complementary strengths for detecting condition-dependent organelle phenotypes.

Pipelines well suited for lipid droplet detection are not necessarily optimal for mitochondrial segmentation or network analysis, highlighting the limitations of universal workflows for structurally distinct organelles. Instead, pipeline selection should be guided by the biological question, organelle of interest, imaging modality, and practical considerations such as automation level, execution time, and ease of use. For lipid droplet analysis, ilastik yielded segmentation that was more consistent across modalities, while for fluorescence-based mitochondrial imaging several pipelines performed comparably. In contrast, none of the evaluated approaches yielded fully robust mitochondrial segmentation for holotomographic images, underscoring current limitations in label-free mitochondrial analysis.

A limitation of this study is that all analyses were performed on two-dimensional representations derived from three-dimensional image stacks. Both lipid droplets and mitochondrial networks are inherently three-dimensional, and projection or slice selection can introduce artifacts, merge structures, or alter apparent connectivity. Fully 3D segmentation and network analysis may reduce some imaging modality- and pipeline-dependent effects observed here but remain less standardized and are not yet broadly supported across all evaluated platforms. Although deep learning performance may improve with retraining or fine-tuning on domain-specific data, our evaluation reflects the use of pretrained models as provided, with no additional training or fine-tuning.

Overall, this work emphasizes the need for careful pipeline selection, standardized downstream analysis, and routine visual validation when performing quantitative organelle analysis (Table 1). By providing reproducible workflows and a systematic comparison across commonly used tools, this study offers practical guidance for selecting appropriate analysis strategies and highlights key considerations for improving reproducibility and interpretability in quantitative cell imaging studies.

## Methods

### Cell culture

The human osteosarcoma cell line TE85 was used as a model for this study. The cells were maintained in high-glucose DMEM containing GlutaMAX™ and sodium pyruvate (Thermo Fisher Scientific, cat. no. 31966-047), supplemented with 10% fetal bovine serum (FBS) and 1% penicillin–streptomycin (hereafter referred to as complete medium). The cells were cultured at 37 °C in a humidified atmosphere with 5% CO₂.

For all experiments, the cells were seeded at a density of 2.6 × 10³ cells/cm² in Cellvis 24-well imaging bottom plates (P24-1.5P) and cultured for 48 h prior to imaging, reaching approximately 60–70% confluency at the time of imaging.

For staining experiments, cells were transferred to serum-free DMEM supplemented with 1% penicillin–streptomycin, as recommended by the manufacturer.

### Fluorescent staining

Prior to staining, the cells were washed with phosphate-buffered saline (PBS). Mitochondria were labeled by incubating cells with MitoTracker™ Deep Red FM (Thermo Fisher Scientific, cat. no. M22426) at 25 nM for 30 min at 37 °C in serum-free DMEM. The cells were then washed with PBS and incubated with BODIPY 493/503 (Thermo Fisher Scientific, cat. no. D3922) at 1 µM to label lipid dropletsfor 20 min in serum-free DMEM.

After staining, cells were washed with PBS, returned to complete medium, and immediately imaged for downstream analysis.

### Image acquisition

Live-cell imaging was performed using an HT-X1™ holotomographic microscope (TomoCube), based on optical diffraction tomography for label-free three-dimensional refractive index imaging^27,54,55^. The cells were maintained at 37 °C a humidified atmosphere with 5% CO₂ during image acquisition using the microscope’s environmental chamber. The refractive index of the surrounding medium was set to RI = 1.337.

Holotomographic images were acquired as three-dimensional z-stacks with a z-step size of 0.78 µm. Fluorescence acquisition settings were optimized prior to imaging and kept constant across all samples. Excitation intensity and exposure time were set to 50% and 200 ms for BODIPY and 10% and 100 ms for MitoTracker, respectively. Fluorescence images were acquired as three-dimensional z-stacks with a z-step size of 0.78 µm.

Holotomographic and fluorescence image stacks were acquired sequentially for each field of view. In total, 104 images were taken across 8 biological replicates.

### Holotomographic image preprocessing

For holotomographic data, three-dimensional z-stacks were converted into two-dimensional representation. For lipid droplet analysis, a maximum intensity projection (MIP) was generated. For mitochondrial analysis, a single best-focus slice was selected using a Laplacian-variance focus metric (implemented as the slice with the highest Laplacian variance across the z-stack).

After z-stack reduction, images were intensity-normalized using min–max scaling (subtracting the minimum intensity and dividing by the maximum intensity) and exported in both 16-bit TIFF and 8-bit PNG formats for downstream analysis.

### Fluorescence image preprocessing

For fluorescence data, three-dimensional z-stacks were converted into two-dimensional images using maximum intensity projection (MIP) for all analyzed structures. MIP images were converted to floating-point representation and intensity-normalized using min–max scaling prior to export as 8-bit TIFF images for downstream analysis, using custom Python scripts.

For lipid droplet fluorescence images, no additional preprocessing beyond MIP generation and intensity rescaling for export was applied.

For mitochondrial fluorescence images, additional preprocessing was applied to enhance filamentous structures prior to segmentation. Specifically, MIP images were subjected to Gaussian background estimation and subtraction, followed by Gaussian smoothing, contrast-limited adaptive histogram equalization (CLAHE), and gamma correction. Both the raw MIP images and the preprocessed images were saved and used for downstream segmentation and quantitative analysis.

### Image analysis and segmentation

Segmentation and quantitative analysis were performed using multiple image analysis software tools, including Fiji (ImageJ), custom Python scripts, ilastik, CellProfiler, TomoAnalysis, MitoSegNet/MitoA, MitoAnalyzer, and MoDL. All analyses were applied to the same preprocessed images to enable direct comparison across methods.

Analyses were performed using Fiji (ImageJ, version 1.54p) ^32,33^, Python (version 3.13.7), ilastik (version 1.4.1.post1) ^37^, CellProfiler (version 4.2.8)^31^, TomoAnalysis (version 2.1.10), MitoSegNet^35^, MitoA^35^, MitoAnalyzer^29^, and MoDL^36^.

Whenever supported by the software, segmentation and quantitative feature extraction were performed within the same environment; otherwise, binary masks were exported and analyzed using identical downstream workflows to ensure comparability across pipelines. Except where required by the software design (e.g. supervised training in ilastik), parameters were not optimized on a per-image or per-condition basis. Representative images used for threshold determination or classifier training were selected to capture intensity and structural variability across the dataset.

Detailed pipeline schematics and parameter settings for each analysis workflow are provided in Supplementary Figures S1–S4. All Fiji macros and Python scripts used for segmentation and quantitative analysis are provided with the manuscript.

#### Fiji (ImageJ)

Fiji-based analyses were performed on preprocessed two-dimensional images using custom batch macros, with parameters kept consistent within each analysis pipeline. Thresholding strategies depended on the cellular structure and imaging modality, but all thresholding was implemented using Fiji’s *SetThreshold* function with default parameters. For holotomographic lipid droplet analysis, image-specific thresholding was applied to account for intensity variability between images (Supplementary Figure S1A). For all other analyses (fluorescence lipid droplets and mitochondria), fixed global thresholds were used, with threshold values determined on a subset of representative images and applied uniformly to all corresponding images (Supplementary Figures S2A, S3A, S4A).

For lipid droplet analysis, binary masks were refined using watershed segmentation, and individual droplets were quantified using the *Analyze Particles* function to extract size and shape descriptors. For mitochondrial analysis, binary masks were quantified using *Analyze Particles* without watershed segmentation to extract morphological features, and subsequently skeletonized using the *Skeletonize* function. Skeletonized images were analyzed with *the Analyze Skeleton* plugin to quantify mitochondrial network topology features (Supplementary Figures S1A, S2A, S3A, S4A).

#### Python

Python-based analyses were performed using custom scripts applied to the same preprocessed images. Numerical operations and data handling were performed using NumPy^56^ and pandas^57^. Image I/O, connected-component labeling, region property extraction, and skeletonization were implemented using scikit-image^58^.

For holotomographic lipid droplet analysis, binary masks were generated by applying a fixed global intensity threshold derived from the mean threshold used in the Fiji-based analysis (Supplementary Figure S1B), followed by connected-component labeling. For each detected object, size and shape descriptors were extracted using region properties, including area, perimeter, major and minor axis lengths, eccentricity, and solidity. Fiji-like shape metrics were additionally calculated, including circularity 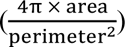, aspect ratio 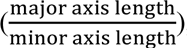, and roundness 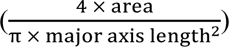. Measurements were converted into physical units using the imaging pixel size.

For fluorescence analyses, the same region-based feature extraction workflow was applied to binary masks generated using fixed global thresholding, consistent with the Fiji-based analysis. Object-level measurements were aggregated per image by calculating the mean and standard deviation, and results were exported as per-object and per-image summary tables.

For mitochondrial network analysis in Python, binary masks were skeletonized using the skeletonize function from scikit-image. Skeleton topology metrics were extracted using graph-based analysis, with skeleton graphs constructed using skan^59^ and network metrics computed using NetworkX^60^. Quantified features included the number of branches, junctions, and endpoints, as well as branch length statistics (total, mean, and maximum branch length per image). All outputs were exported as per-object/per-branch tables and per-image summary files (Supplementary Figures S2B, S3B, S4B).

#### ilastik

Segmentation using ilastik was performed via supervised pixel classification, with separate classifiers trained for each cellular structure and imaging modality. Pixels were manually labeled into two classes foreground (lipid droplets or mitochondria) and background. For lipid droplet segmentation, classifiers were trained using 10 manually annotated images for fluorescence data and 10 manually annotated images for holotomographic data. For mitochondrial segmentation, classifiers were trained using 10 manually annotated fluorescence images, whereas a larger training set of 54 manually annotated holotomographic images was used to capture the increased structural complexity of mitochondrial networks in holotomographic data (Supplementary Figures S2C and S4C). All available ilastik image features were selected during training. Binary segmentation masks generated by ilastik were exported for downstream analysis. Subsequent quantitative feature extraction and mitochondrial network analysis were performed using the same Fiji- and Python-based workflows described above.

#### CellProfiler

CellProfiler was used for fluorescence-based lipid droplet and mitochondria segmentation and quantitative analysis using a predefined analysis pipeline. Images were first intensity-normalized using the *RescaleIntensity* module and enhanced using *EnhanceOrSuppressFeatures* to improve lipid droplet contrast. Lipid droplets were identified using *IdentifyPrimaryObjects* with global thresholding.

Object-level lipid droplet size and shape features were quantified directly within CellProfiler using *MeasureObjectSizeShape* and *MeasureObjectIntensity*. Segmented objects and measurement tables were exported using *SaveImages* and *ExportToSpreadsheet*.

For mitochondrial analysis, binary masks generated by CellProfiler were exported and used exclusively for downstream mitochondrial network (skeleton-based) analysis, which was performed using the Fiji-based workflows described above (Supplementary Figures S2D, S4D).

#### TomoAnalysis

TomoAnalysis was used for lipid droplet segmentation from holotomographic images using the built-in two-dimensional lipid droplet analysis pipeline, following the workflow recommended by the software developers. Segmentation was performed directly within TomoAnalysis on two-dimensional representations of holotomographic data. Lipid droplet detection and quantitative measurements were performed directly within TomoAnalysis.

Only lipid droplet number and area measurements were available and included in the comparative analysis.

#### MitoSegNet

Mitochondrial segmentation was performed using the MitoS segmentation tool, which implements the pretrained deep learning model MitoSegNet, following the workflow recommended by the developers.

Segmentation was applied to two-dimensional fluorescence images converted to 8-bit format, as required by the MitoS tool.

The pretrained MitoSegNet model was used in Basic Mode without fine-tuning. No size exclusion was applied to ensure consistency across pipelines. Binary mitochondrial segmentation masks were generated directly by the model.

Binary masks were subsequently analyzed using the MitoA analyzer tool, as recommended by the developers, to extract mitochondrial morphological and network-related features. These features were included alongside results obtained from other analysis pipelines for comparative analysis.

#### MoDL

Mitochondrial segmentation using MoDL was performed using the default pretrained model and parameter settings recommended by the developers. No additional parameter tuning or retraining was performed.

Binary mitochondrial masks generated by MoDL were exported and used for downstream quantitative analysis, including morphological and network-level feature extraction, following the same analysis strategy applied to other segmentation methods.

#### MitoAnalyzer

MitoAnalyzer was applied to binary mitochondrial masks generated using the same intensity thresholding strategy as the Fiji-based mitochondrial segmentation. Mitochondrial morphological and network-related features were extracted using the built-in analysis routines. Analysis was performed in per-cell mode without supplying cell masks or regions of interest (Mask Channel = None), such that the full field of view was treated as a single region and features were reported on a per-image basis. Because the MitoAnalyzer plugin does not support batch processing with externally defined thresholds, a custom script was implemented to enable automated analysis across all images using identical input mask.

### Statistical analyses

Quantitative features were first computed at the individual object level and subsequently aggregated per image by calculating either the mean or the standard deviation, depending on the analysis. Statistical comparisons between image analysis pipelines were performed on these per-image summary values. Because the same images were analyzed using multiple pipelines, measurements were treated as repeated measures. Overall differences across pipelines were assessed using repeated-measures one-way ANOVA with Geisser–Greenhouse correction to account for violations of sphericity. When the ANOVA indicated a significant effect, post hoc pairwise comparisons between pipelines were performed using paired tests with appropriate multiple-comparison correction. A significance threshold of p < 0.05 was used throughout the study. Selected statistical comparisons are indicated directly on the plots, while complete statistical results for all pairwise comparisons are reported in the Supplementary Tables. Selected statistical comparisons are indicated directly on the plots, while complete statistical results for all pairwise comparisons are reported in the Supplementary Tables.

For comparisons between two independent experimental conditions (e.g., control versus serum starvation), statistical significance was assessed using Welch’s t-test on per-image summary values. A significance threshold of p < 0.05 was used throughout the study. Statistical comparisons are indicated directly on the plots.

### Pairwise comparison of mitochondrial skeleton overlap

Agreement between mitochondrial skeletons generated by different analysis pipelines was quantified using the Dice coefficient and a relaxed Dice coefficient. The Dice coefficient measures how much two binary masks overlap and is defined as:

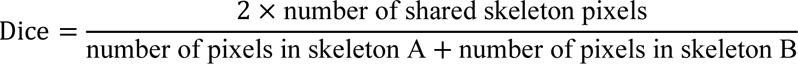

where skeleton A and skeleton B correspond to the binary skeleton masks produced by two different analysis pipelines.

To account for small spatial offsets between skeleton representations, a relaxed Dice coefficient was additionally computed. For this metric, each skeleton was symmetrically dilated by 2 pixels, and overlap was calculated between each original skeleton and the dilated version of the other. This relaxed measure reduces sensitivity to minor positional differences while preserving sensitivity to overall network agreement.

Pairwise Dice and relaxed Dice values were assembled into symmetric similarity matrices, with diagonal values set to 1. These matrices were visualized as heatmaps to summarize agreement between analysis pipelines.

### Oleic acid treatment and lipid droplet quantification by holotomography

To reduce basal lipid availability prior to treatment, cells were starved for 24 h in serum-free high-glucose DMEM containing GlutaMAX™ and sodium pyruvate (Thermo Fisher Scientific, cat. no. 31966-047) supplemented with 1% penicillin–streptomycin (PS). Oleic acid treatment was then performed in the same serum-free medium, with the addition of 1% BSA as a fatty-acid carrier. Oleic acid was added to achieve a final concentration of 2 mM, and cells were incubated 24 h prior to imaging.

For identification and counting, cells were stained with Hoechst 33342 (Bio-Techne, cat. no. 5117/50) at a final concentration of 37.5 µM for 20 min, after which imaging was performed immediately.

Lipid droplet segmentation masks were generated using the holotomography lipid droplet Python analysis script, applying the exact same segmentation threshold parameters as in the previous analysis to ensure comparability across experiments. Per-image summary metrics were computed.

To account for differences in cell density across fields of view, the number of lipid droplets was normalized by the number of cells per image (cell counts derived from Hoechst-stained nuclei). In contrast, droplet morphology metrics (area, aspect ratio, roundness) were calculated at the object level and summarized per image without normalization to cell number.

Experiments were performed across 5 biological replicates, with a total of 50 iamges analyzed per condition.

### Serum starvation and mitochondrial morphology quantification

To induce metabolic stress prior to imaging, cells in the starvation condition were incubated for 24 h in serum-free high-glucose DMEM supplemented with GlutaMAX™, sodium pyruvate, and 1% penicillin–streptomycin. Control cells were maintained in complete growth medium for the same duration.

For simultaneous nuclear and mitochondrial labeling, cells were co-stained with Hoechst 33342 (Bio-Techne, cat. no. 5117/50) at a final concentration of 37.5 µM and MitoTracker™ Deep Red FM (Thermo Fisher Scientific, cat. no. M22426) at a final concentration of 25 nM in serum-free DMEM supplemented with 1% penicillin–streptomycin. Prior to staining, cells were washed with phosphate-buffered saline (PBS). Cells were incubated for 30 min at 37 °C, after which the staining medium was replaced with complete growth medium and imaging was performed immediately without an additional wash step.

Mitochondrial segmentation masks were generated using the previously trained ilastik classifier applied uniformly across all images. Morphometric and network features were extracted from the resulting masks, including mitochondrial object count, area, perimeter, and aspect ratio, as well as skeleton-based network metrics (total branches, branch junctions, branch endpoints, total branch length, mean branch length, and maximum branch length). Per-image summary metrics were computed for downstream analysis.

To account for differences in cell density across fields of view, mitochondrial network features including mitochondrial object number, branch number, junction number, endpoint number, and total branch length, were normalized to the number of cells per image, obtained from nuclear segmentation. In contrast, mitochondrial morphology metrics (area, perimeter, aspect ratio, mean branch length, and maximum branch length) were computed at the object level and summarized per image without normalization to cell number. Network topology metrics were calculated from skeletonized mitochondrial masks and reported per image. The same segmentation and feature-extraction parameters were applied across all datasets to ensure consistency between starved and control conditions.

Experiments were performed across 5 biological replicates, with a total of 50 images analyzed per condition.

## ASSOCIATED CONTENT

Supplementary information can be found in *Supplementary Information.docx*

Numerical Values can be found in *Numerical Values.xlsx*

Code can be found in https://github.com/chloedaul/organelle-quantification.git

## AUTHOR INFORMATION

### Corresponding Author

* Shukry J. Habib

## Author contribution

The manuscript was written through contributions of all authors. All authors have given approval to the final version of the manuscript.

**Chloe Daul:** Data curation, Formal analysis, Validation, Investigation, Visualization, Methodology, Writing – original draft, Writing – review and editing.

**Pierre Tournier:** Conceptualization, Investigation, Methodology, Supervision, Project administration, Writing – review and editing.

**Shukry J. Habib:** Conceptualization, Resources, Supervision, Funding acquisition, Project administration, Writing – review and editing.

## ACKNOWLEDGMENT

The authors acknowledge the University of Lausanne for its financial support.

