## Supplementary Information for "Benchmarking Machine Learning and Automated Image Analysis for Organelle Quantification"

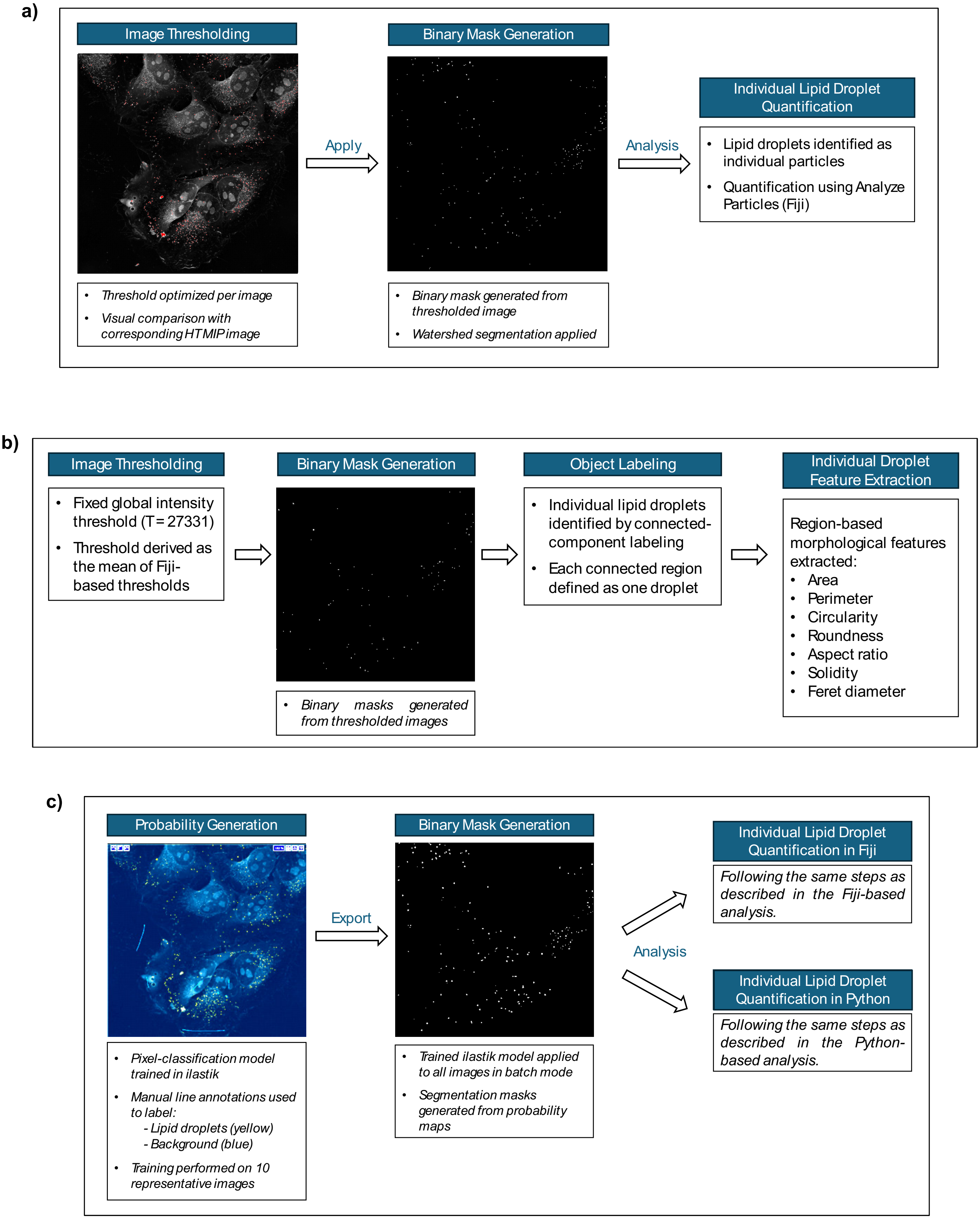

**Supplementary Figure S1**. Overview of image analysis pipelines for lipid droplet segmentation and quantification in holotomographic images. Panels (a-c) summarize the different analysis pipelines applied to label-free holotomographic (HT) images for lipid droplet segmentation and quantification. (a) Fiji pipeline applied to holotomographic images. Three-dimensional holotomographic stacks were converted into two-dimensional representations using maximum intensity projection (MIP). Lipid droplets were segmented using image-specific intensity thresholding to account for variability in refractive index contrast across images, followed by binary mask generation and watershed segmentation. Individual lipid droplets were identified and quantified using the Analyze Particles tool to extract size and shape descriptors. (b) Python-based pipeline for holotomographic images. Maximum intensity projections were segmented using a fixed global intensity threshold derived from the Fiji-based analysis. Binary masks were generated and lipid droplets were identified via connected-component labeling. Morphological features, including area, perimeter, circularity, aspect ratio, and roundness, were automatically extracted for each detected droplet. (c) ilastik pipeline for holotomographic images. Supervised pixel classification was performed using manually annotated training images to generate lipid droplet probability maps. Probability maps were converted into binary segmentation masks and exported for downstream analysis. Lipid droplet quantification was subsequently performed using the same Fiji- or Python-based workflows to ensure comparability across pipelines.

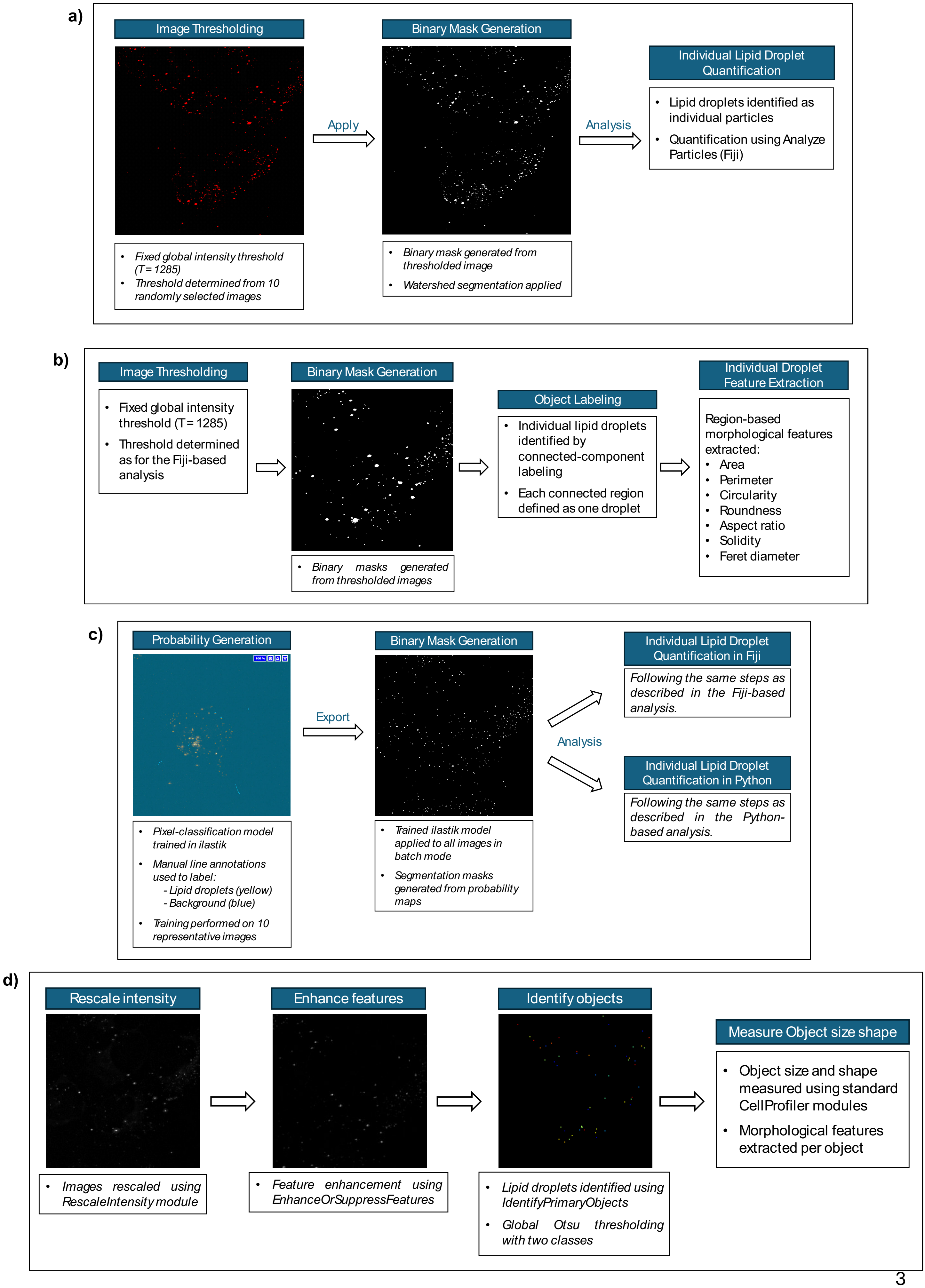
a)
Image Thresholding
Apply
Binary Mask Generation
Analysis
Individual Lipid Droplet Quantification
Lipid droplets identified as individual particles
Quantification using Analyze Particles (Fiji)
Fixed global intensity threshold (T = 1285)
Threshold determined from 10 randomly selected images
Binary mask generated from thresholded image
Watershed segmentation applied
b)
Image Thresholding
Fixed global intensity threshold (T = 1285)
Threshold determined as for the Fiji-based analysis
Binary Mask Generation
Binary masks generated from thresholded images
Object Labeling
Individual lipid droplets identified by connected-component labeling
Each connected region defined as one droplet
Individual Droplet Feature Extraction
Region-based morphological features extracted:
Area
Perimeter
Circularity
Roundness
Aspect ratio
Solidity
Feret diameter
c)
Probability Generation
Export
Binary Mask Generation
Analysis
Individual Lipid Droplet Quantification in Fiji
Following the same steps as described in the Fiji-based analysis.
Individual Lipid Droplet Quantification in Python
Following the same steps as described in the Python-based analysis.
Pixel-classification model trained in ilastik
Manual line annotations used to label:
- Lipid droplets (yellow)
- Background (blue)
Training performed on 10 representative images
Trained ilastik model applied to all images in batch mode
Segmentation masks generated from probability maps
d)
Rescale intensity
Enhance features
Identify objects
Measure Object size shape
Object size and shape measured using standard CellProfiler modules
Morphological features extracted per object
Images rescaled using RescaleIntensity module
Feature enhancement using EnhanceOrSuppressFeatures
Lipid droplets identified using IdentifyPrimaryObjects
Global Otsu thresholding with two classes

**Supplementary Figure S2**. Overview of image analysis pipelines for lipid droplet segmentation and quantification in fluorescence images. Panels (a-d) summarize the different analysis pipelines applied to fluorescence microscopy images for lipid droplet analysis. (a) Fiji pipeline applied to fluorescence images. Three-dimensional fluorescence z-stacks were converted into two-dimensional maximum intensity projections. Lipid droplets were segmented using a fixed global intensity threshold defined from a subset of representative images, followed by binary mask generation and watershed segmentation. Individual lipid droplets were quantified using Analyze Particles to extract size and shape features. (b) Python-based pipeline for fluorescence images. Fluorescence projections were segmented using the same global thresholding parameters as the Fiji-based workflow. Binary masks were generated and lipid droplets were identified using connected-component labeling. Morphological features were automatically extracted using region-based measurements implemented in Python. (c) ilastik pipeline for fluorescence images. Supervised pixel classification was used to generate lipid droplet probability maps from fluorescence images. Binary segmentation masks derived from these probability maps were exported and analyzed using standardized Fiji- or Python-based quantification workflows. (d) CellProfiler pipeline for fluorescence images. Fluorescence images were intensity-normalized and enhanced using standard CellProfiler modules. Lipid droplets were identified using global thresholding in the IdentifyPrimaryObjects module. Object-level size and shape measurements were extracted directly within CellProfiler and exported for comparative analysis.

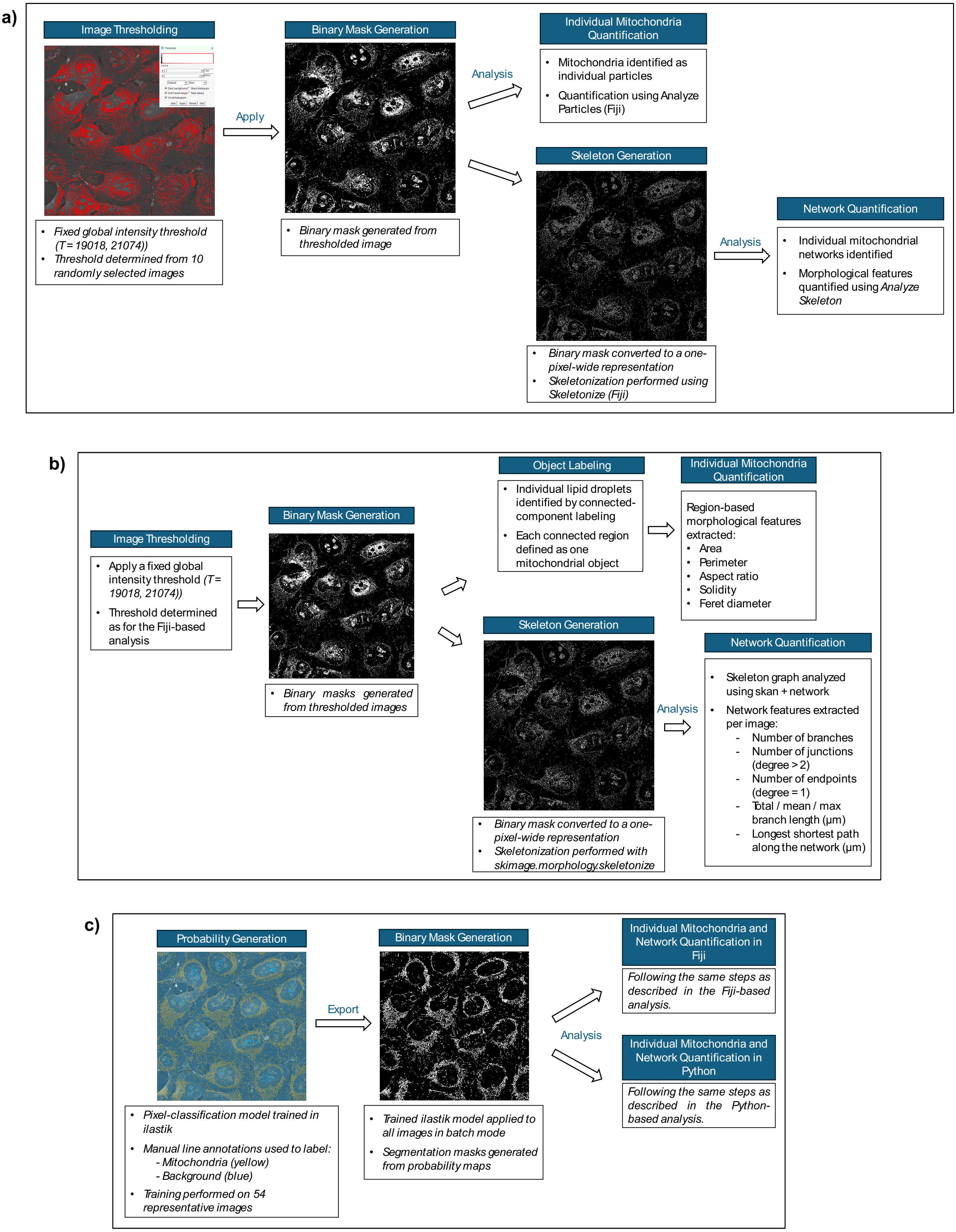

a)
Image Thresholding
Apply
Binary Mask Generation
Analysis
Individual Mitochondria Quantification
• Mitochondria identified as individual particles
• Quantification using Analyze Particles (Fiji)
Skeleton Generation
• Fixed global intensity threshold (T = 19018, 21074))
• Threshold determined from 10 randomly selected images
• Binary mask generated from thresholded image
Analysis
Network Quantification
• Individual mitochondrial networks identified
• Morphological features quantified using Analyze Skeleton
• Binary mask converted to a one-pixel-wide representation
• Skeletonization performed using Skeletonize (Fiji)
b)
Object Labeling
• Individual lipid droplets identified by connected-component labeling
• Each connected region defined as one mitochondrial object
Individual Mitochondria Quantification
Region-based morphological features extracted:
• Area
• Perimeter
• Aspect ratio
• Solidity
• Feret diameter
Image Thresholding
• Apply a fixed global intensity threshold (T = 19018, 21074))
• Threshold determined as for the Fiji-based analysis
Binary Mask Generation
• Binary masks generated from thresholded images
Skeleton Generation
Analysis
Network Quantification
• Skeleton graph analyzed using skan + network
• Network features extracted per image:
- Number of branches
- Number of junctions (degree > 2)
- Number of endpoints (degree = 1)
- Total / mean / max branch length (µm)
- Longest shortest path along the network (µm)
• Binary mask converted to a one-pixel-wide representation
• Skeletonization performed with skimage.morphology.skeletonize
c)
Probability Generation
Export
Binary Mask Generation
Individual Mitochondria and Network Quantification in Fiji
Following the same steps as described in the Fiji-based analysis.
Analysis
Individual Mitochondria and Network Quantification in Python
Following the same steps as described in the Python-based analysis.
• Pixel-classification model trained in ilastik
• Manual line annotations used to label:
- Mitochondria (yellow)
- Background (blue)
• Training performed on 54 representative images
• Trained ilastik model applied to all images in batch mode
• Segmentation masks generated from probability maps

**Supplementary Figure S3.** Overview of image analysis pipelines for mitochondrial segmentation and network quantification in holotomographic images. Panels (a-c) summarize the different analysis pipelines applied to label-free holotomographic (HT) images for mitochondrial segmentation and network quantification. (a) Fiji pipeline applied to holotomographic images. Three-dimensional holotomographic stacks were reduced to a single best-focus slice using a Laplacian-variance focus metric. Mitochondria were segmented using fixed global intensity thresholding defined from a subset of representative images, followed by binary mask generation. Skeletons were generated using the Fiji *Skeletonize* function, and mitochondrial network features were quantified using the *Analyze Skeleton* plugin. (b) Python-based pipeline for holotomographic images. Best-focus slices were segmented using the same global thresholding strategy as in the Fiji pipeline. Binary masks were skeletonized using scikit-image, and mitochondrial network graphs were constructed using *skan* and *NetworkX* to extract branch counts, junctions, endpoints, and branch length statistics. (c) ilastik pipeline for holotomographic images. Supervised pixel classification was performed using manually annotated training images to generate mitochondrial probability maps. Probability maps were converted into binary segmentation masks, which were subsequently analyzed using the same Fiji- or Python-based skeletonization and network quantification workflows described above.

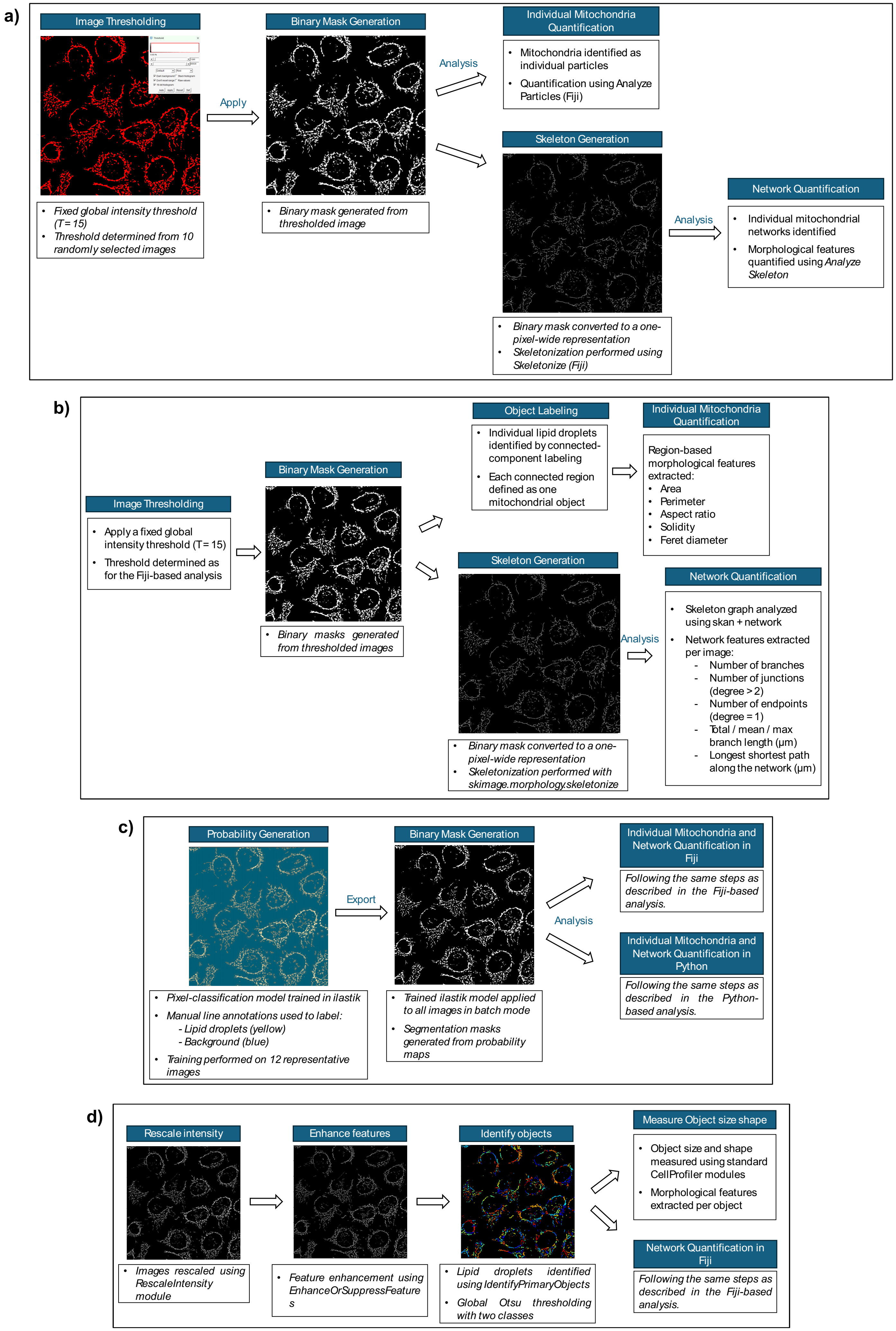
a)
Image Thresholding
Apply
Fixed global intensity threshold (T= 15)
Threshold determined from 10 randomly selected images
Binary Mask Generation
Binary mask generated from thresholded image
Analysis
Individual Mitochondria Quantification
Mitochondria identified as individual particles
Quantification using Analyze Particles (Fiji)
Skeleton Generation
Binary mask converted to a one-pixel-wide representation
Skeletonization performed using Skeletonize (Fiji)
Analysis
Network Quantification
Individual mitochondrial networks identified
Morphological features quantified using Analyze Skeleton
b)
Image Thresholding
Apply a fixed global intensity threshold (T= 15)
Threshold determined as for the Fiji-based analysis
Binary Mask Generation
Binary masks generated from thresholded images
Object Labeling
Individual lipid droplets identified by connected-component labeling
Each connected region defined as one mitochondrial object
Individual Mitochondria Quantification
Region-based morphological features extracted:
Area
Perimeter
Aspect ratio
Solidity
Feret diameter
Skeleton Generation
Binary mask converted to a one-pixel-wide representation
Skeletonization performed with skimage.morphology.skeletonize
Analysis
Network Quantification
Skeleton graph analyzed using skan + network
Network features extracted per image:
- Number of branches
- Number of junctions (degree > 2)
- Number of endpoints (degree = 1)
- Total / mean / max branch length (µm)
- Longest shortest path along the network (µm)
c)
Probability Generation
Pixel-classification model trained in ilastik
Manual line annotations used to label:
- Lipid droplets (yellow)
- Background (blue)
Training performed on 12 representative images
Export
Binary Mask Generation
Trained ilastik model applied to all images in batch mode
Segmentation masks generated from probability maps
Analysis
Individual Mitochondria and Network Quantification in Fiji
Following the same steps as described in the Fiji-based analysis.
Individual Mitochondria and Network Quantification in Python
Following the same steps as described in the Python-based analysis.
d)
Rescale intensity
Images rescaled using RescaleIntensity module
Enhance features
Feature enhancement using EnhanceOrSuppressFeatures
Identify objects
Lipid droplets identified using IdentifyPrimaryObjects
Global Otsu thresholding with two classes
Measure Object size shape
Object size and shape measured using standard CellProfiler modules
Morphological features extracted per object
Network Quantification in Fiji
Following the same steps as described in the Fiji-based analysis.

**Supplementary Figure S4.** Overview of image analysis pipelines for mitochondrial segmentation and network quantification in fluorescence images. Panels (a-d) summarize the analysis pipelines applied to fluorescence microscopy images for mitochondrial segmentation and network analysis. (a) Fiji pipeline applied to fluorescence images. Three-dimensional fluorescence z-stacks were converted into two-dimensional maximum intensity projections. Images were preprocessed to enhance filamentous mitochondrial structures, followed by global intensity thresholding and binary mask generation. Skeletonization and network quantification were performed using the Fiji *Skeletonize* and *Analyze Skeleton* tools. (b) Python-based pipeline for fluorescence images. Preprocessed fluorescence projections were segmented using global thresholding parameters matched to the Fiji-based workflow. Binary masks were skeletonized using scikit-image, and mitochondrial network features were extracted using graph-based analysis implemented with *skan* and *NetworkX*. (c) *ilastik pipeline for fluorescence images.* Supervised pixel classification was used to generate mitochondrial probability maps from fluorescence images. Binary segmentation masks derived from probability maps were subsequently analyzed using the same Fiji- or Python-based skeletonization and network quantification workflows. (d) CellProfiler pipeline for fluorescence images. Fluorescence images were intensity-rescaled and enhanced using standard CellProfiler modules. Mitochondria were identified using global thresholding in the *IdentifyPrimaryObjects* module. Binary masks were exported and subjected to downstream skeletonization and network analysis using Fiji-based workflows.

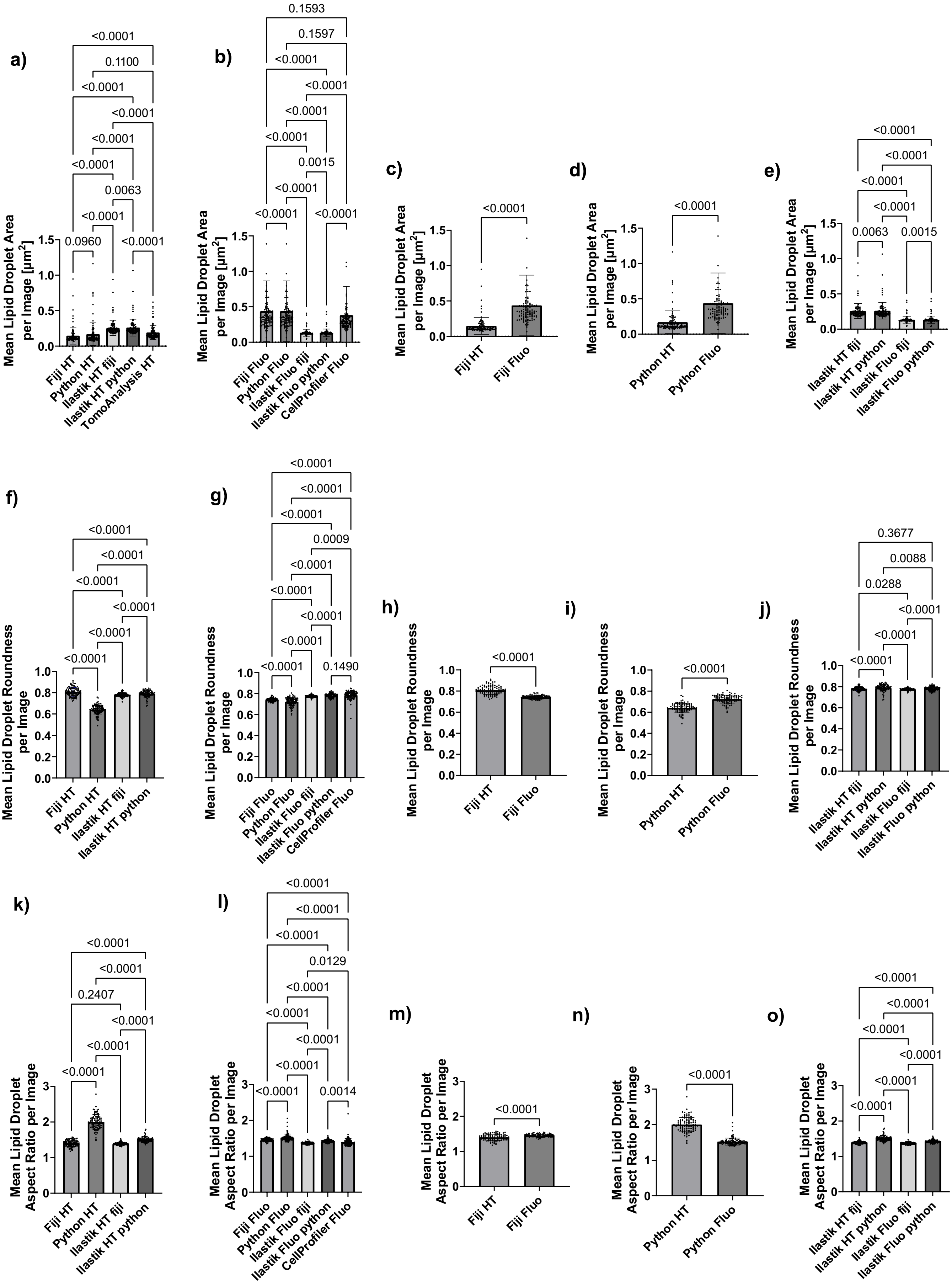

a)
Mean Lipid Droplet Area per Image [µm²]
Fiji HT
Python HT
Ilastik HT fiji
Ilastik HT python
TomoAnalysis HT
<0.0001
0.1100
<0.0001
<0.0001
<0.0001
<0.0001
0.0063
<0.0001
0.0960
<0.0001
b)
Mean Lipid Droplet Area per Image [µm²]
Fiji Fluo
Python Fluo
Ilastik Fluo fiji
Ilastik Fluo python
CellProfiler Fluo
0.1593
0.1597
<0.0001
<0.0001
<0.0001
<0.0001
0.0015
<0.0001
<0.0001
<0.0001
c)
Mean Lipid Droplet Area per Image [µm²]
Fiji HT
Fiji Fluo
<0.0001
d)
Mean Lipid Droplet Area per Image [µm²]
Python HT
Python Fluo
<0.0001
e)
Mean Lipid Droplet Area per Image [µm²]
Ilastik HT fiji
Ilastik HT python
Ilastik Fluo fiji
Ilastik Fluo python
<0.0001
<0.0001
<0.0001
<0.0001
0.0063
0.0015
f)
Mean Lipid Droplet Roundness per Image
Fiji HT
Python HT
Ilastik HT fiji
Ilastik HT python
<0.0001
<0.0001
<0.0001
<0.0001
<0.0001
<0.0001
g)
Mean Lipid Droplet Roundness per Image
Fiji Fluo
Python Fluo
Ilastik Fluo fiji
Ilastik Fluo python
CellProfiler Fluo
<0.0001
<0.0001
<0.0001
0.0009
<0.0001
<0.0001
<0.0001
<0.0001
<0.0001
0.1490
h)
Mean Lipid Droplet Roundness per Image
Fiji HT
Fiji Fluo
<0.0001
i)
Mean Lipid Droplet Roundness per Image
Python HT
Python Fluo
<0.0001
j)
Mean Lipid Droplet Roundness per Image
Ilastik HT fiji
Ilastik HT python
Ilastik Fluo fiji
Ilastik Fluo python
0.3677
0.0088
0.0288
<0.0001
<0.0001
<0.0001
k)
Mean Lipid Droplet Aspect Ratio per Image
Fiji HT
Python HT
Ilastik HT fiji
Ilastik HT python
<0.0001
<0.0001
0.2407
<0.0001
<0.0001
<0.0001
l)
Mean Lipid Droplet Aspect Ratio per Image
Fiji Fluo
Python Fluo
Ilastik Fluo fiji
Ilastik Fluo python
CellProfiler Fluo
<0.0001
<0.0001
<0.0001
0.0129
<0.0001
<0.0001
<0.0001
<0.0001
<0.0001
0.0014
m)
Mean Lipid Droplet Aspect Ratio per Image
Fiji HT
Fiji Fluo
<0.0001
n)
Mean Lipid Droplet Aspect Ratio per Image
Python HT
Python Fluo
<0.0001
o)
Mean Lipid Droplet Aspect Ratio per Image
Ilastik HT fiji
Ilastik HT python
Ilastik Fluo fiji
Ilastik Fluo python
<0.0001
<0.0001
<0.0001
<0.0001
<0.0001
<0.0001

**Supplementary Figure S5.** Comparison of mean lipid droplet morphological features across image analysis pipelines. (a-e) Mean lipid droplet area per image. (a) Comparison across analysis pipelines for holotomographic (HT) images. (b) Comparison across analysis pipelines for fluorescence (Fluo) images. (c) Paired comparison of mean lipid droplet area between holotomographic and fluorescence images using the Fiji-based pipeline. (d) Paired comparison of mean lipid droplet area between holotomographic and fluorescence images using the Python-based pipeline. (e) Comparison of mean lipid droplet area obtained using ilastik-based segmentation across imaging modalities and downstream quantification environments. (f-j) Mean lipid droplet roundness per image. (f) Comparison across analysis pipelines for holotomographic images. (g) Comparison across analysis pipelines for fluorescence images. (h) Paired comparison of mean lipid droplet roundness between holotomographic and fluorescence images using the Fiji-based pipeline. (i) Paired comparison of mean lipid droplet roundness between holotomographic and fluorescence images using the Python-based pipeline. (j) Comparison of mean lipid droplet roundness obtained using ilastik-based segmentation across imaging modalities and downstream quantification environments. (k-o) Mean lipid droplet aspect ratio per image. (k) Comparison across analysis pipelines for holotomographic images. (l) Comparison across analysis pipelines for fluorescence images. (m) Paired comparison of mean lipid droplet aspect ratio between holotomographic and fluorescence images using the Fiji-based pipeline. (n) Paired comparison of mean lipid droplet aspect ratio between holotomographic and fluorescence images using the Python-based pipeline. (o) Comparison of mean lipid droplet aspect ratio obtained using ilastik-based segmentation across imaging modalities and downstream quantification environments. Statistical comparisons between pipelines or conditions are indicated directly above each plot.

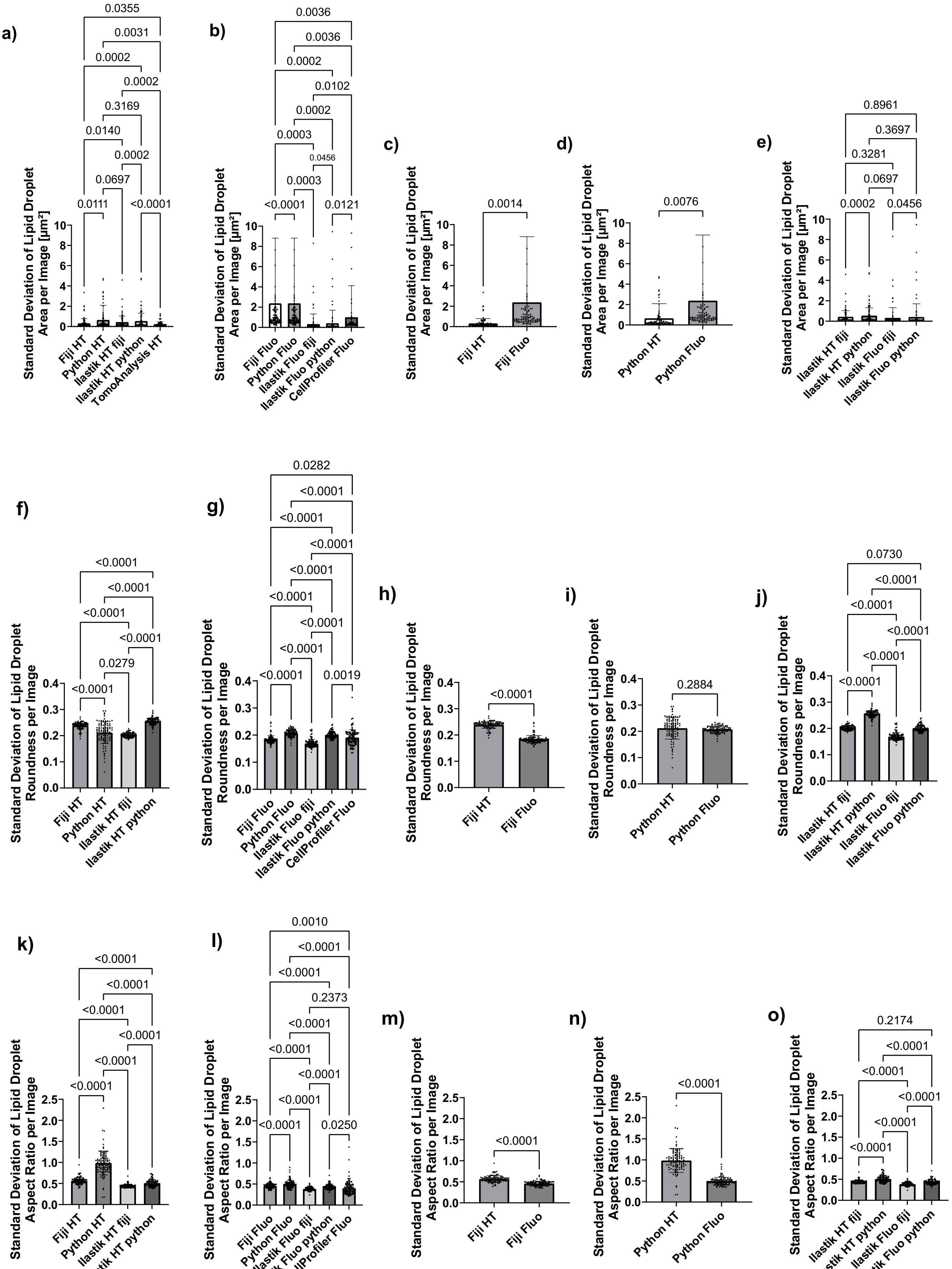

a)
0.0355
0.0031
0.0002
0.0002
0.3169
0.0140
0.0002
0.0697
0.0111
<0.0001
Standard Deviation of Lipid Droplet Area per Image [µm²]
Fiji HT
Python HT
Ilastik HT fiji
Ilastik HT python
TomoAnalysis HT
b)
0.0036
0.0036
0.0002
0.0102
0.0002
0.0003
0.0456
0.0003
<0.0001
0.0121
Standard Deviation of Lipid Droplet Area per Image [µm²]
Fiji Fluo
Python Fluo
Ilastik Fluo fiji
Ilastik Fluo python
CellProfiler Fluo
c)
0.0014
Standard Deviation of Lipid Droplet Area per Image [µm²]
Fiji HT
Fiji Fluo
d)
0.0076
Standard Deviation of Lipid Droplet Area per Image [µm²]
Python HT
Python Fluo
e)
0.8961
0.3697
0.3281
0.0697
0.0002
0.0456
Standard Deviation of Lipid Droplet Area per Image [µm²]
Ilastik HT fiji
Ilastik HT python
Ilastik Fluo fiji
Ilastik Fluo python
f)
<0.0001
<0.0001
<0.0001
<0.0001
0.0279
<0.0001
Standard Deviation of Lipid Droplet Roundness per Image
Fiji HT
Python HT
Ilastik HT fiji
Ilastik HT python
g)
0.0282
<0.0001
<0.0001
<0.0001
<0.0001
<0.0001
<0.0001
<0.0001
<0.0001
0.0019
Standard Deviation of Lipid Droplet Roundness per Image
Fiji Fluo
Python Fluo
Ilastik Fluo fiji
Ilastik Fluo python
CellProfiler Fluo
h)
<0.0001
Standard Deviation of Lipid Droplet Roundness per Image
Fiji HT
Fiji Fluo
i)
0.2884
Standard Deviation of Lipid Droplet Roundness per Image
Python HT
Python Fluo
j)
0.0730
<0.0001
<0.0001
<0.0001
<0.0001
<0.0001
Standard Deviation of Lipid Droplet Roundness per Image
Ilastik HT fiji
Ilastik HT python
Ilastik Fluo fiji
Ilastik Fluo python
k)
<0.0001
<0.0001
<0.0001
<0.0001
<0.0001
<0.0001
Standard Deviation of Lipid Droplet Aspect Ratio per Image
Fiji HT
Python HT
Ilastik HT fiji
Ilastik HT python
l)
0.0010
<0.0001
<0.0001
0.2373
<0.0001
<0.0001
<0.0001
<0.0001
<0.0001
0.0250
Standard Deviation of Lipid Droplet Aspect Ratio per Image
Fiji Fluo
Python Fluo
Ilastik Fluo fiji
Ilastik Fluo python
CellProfiler Fluo
m)
<0.0001
Standard Deviation of Lipid Droplet Aspect Ratio per Image
Fiji HT
Fiji Fluo
n)
<0.0001
Standard Deviation of Lipid Droplet Aspect Ratio per Image
Python HT
Python Fluo
o)
0.2174
<0.0001
<0.0001
<0.0001
<0.0001
<0.0001
Standard Deviation of Lipid Droplet Aspect Ratio per Image
Ilastik HT fiji
Ilastik HT python
Ilastik Fluo fiji
Ilastik Fluo python

**Supplementary Figure S6.** Comparison of lipid droplet feature variability across image analysis pipelines. (a-e) Within-image variability of lipid droplet area, quantified as the standard deviation per image. (a) Comparison across analysis pipelines for holotomographic (HT) images. (b) Comparison across analysis pipelines for fluorescence (Fluo) images. (c) Paired comparison of lipid droplet area variability between holotomographic and fluorescence images using the Fiji-based pipeline. (d) Paired comparison of lipid droplet area variability between holotomographic and fluorescence images using the Python-based pipeline. (e) Comparison of lipid droplet area variability obtained using ilastik-based segmentation across imaging modalities and downstream quantification environments. (f-j) Within-image variability of lipid droplet roundness, quantified as the standard deviation per image. (f) Comparison across analysis pipelines for holotomographic images. (g) Comparison across analysis pipelines for fluorescence images. (h) Paired comparison of lipid droplet roundness variability between holotomographic and fluorescence images using the Fiji-based pipeline. (i) Paired comparison of lipid droplet roundness variability between holotomographic and fluorescence images using the Python-based pipeline. (j) Comparison of lipid droplet roundness variability obtained using ilastik-based segmentation across imaging modalities and downstream quantification environments. (k-o) Within-image variability of lipid droplet aspect ratio, quantified as the standard deviation per image. (k) Comparison across analysis pipelines for holotomographic images. (l) Comparison across analysis pipelines for fluorescence images. (m) Paired comparison of lipid droplet aspect ratio variability between holotomographic and fluorescence images using the Fiji-based pipeline. (n) Paired comparison of lipid droplet aspect ratio variability between holotomographic and fluorescence images using the Python-based pipeline. (o) Comparison of lipid droplet aspect ratio variability obtained using ilastik-based segmentation across imaging modalities and downstream quantification environments. Statistical comparisons between pipelines or conditions are indicated directly above each plot.

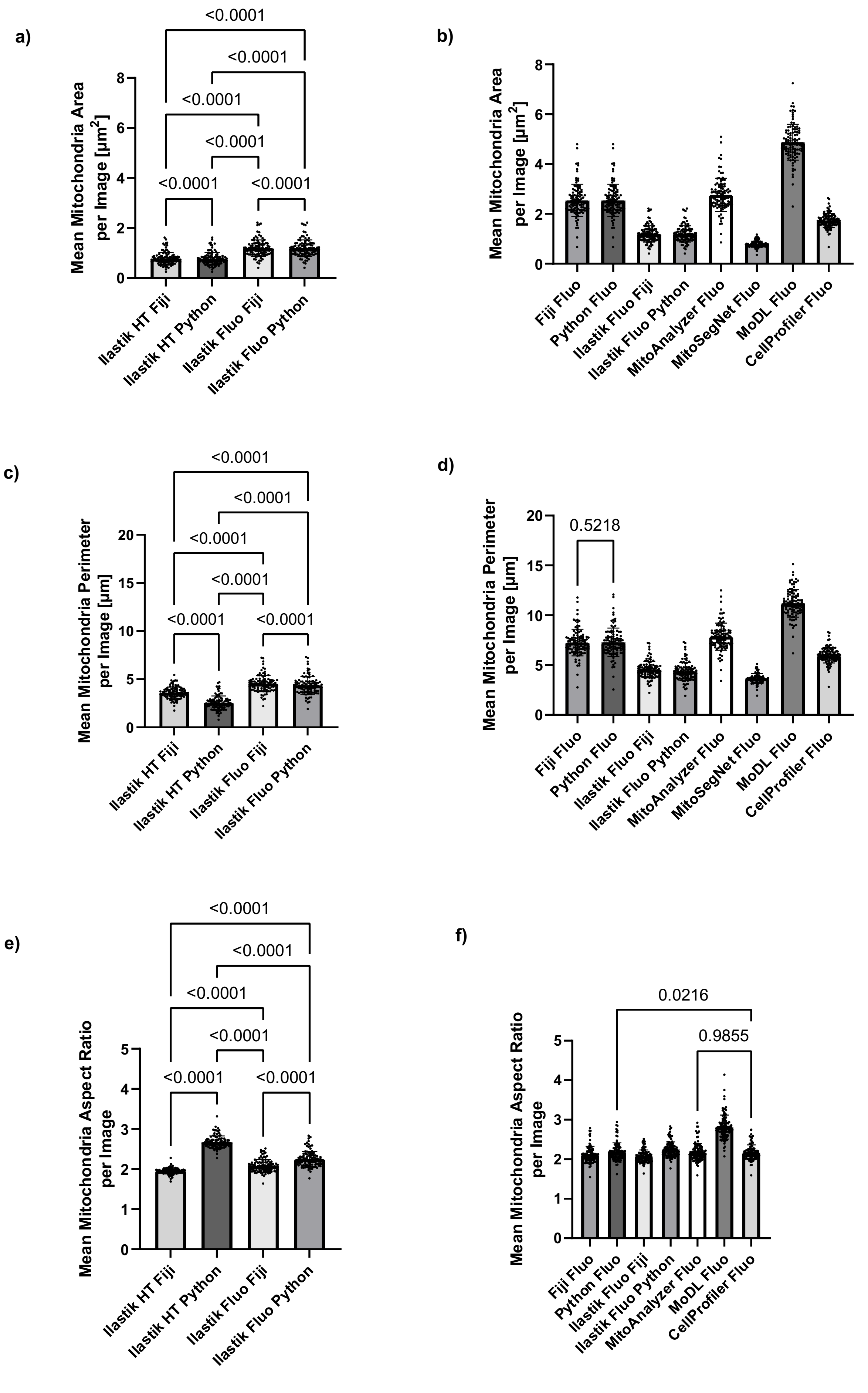
a)
<0.0001
<0.0001
<0.0001
<0.0001
<0.0001
<0.0001
Mean Mitochondria Area per Image [µm2]
Ilastik HT Fiji
Ilastik HT Python
Ilastik Fluo Fiji
Ilastik Fluo Python
b)
Mean Mitochondria Area per Image [µm2]
Fiji Fluo
Python Fluo
Ilastik Fluo Fiji
Ilastik Fluo Python
MitoAnalyzer Fluo
MitoSegNet Fluo
MoDL Fluo
CellProfiler Fluo
c)
<0.0001
<0.0001
<0.0001
<0.0001
<0.0001
<0.0001
Mean Mitochondria Perimeter per Image [µm]
Ilastik HT Fiji
Ilastik HT Python
Ilastik Fluo Fiji
Ilastik Fluo Python
d)
0.5218
Mean Mitochondria Perimeter per Image [µm]
Fiji Fluo
Python Fluo
Ilastik Fluo Fiji
Ilastik Fluo Python
MitoAnalyzer Fluo
MitoSegNet Fluo
MoDL Fluo
CellProfiler Fluo
e)
<0.0001
<0.0001
<0.0001
<0.0001
<0.0001
<0.0001
Mean Mitochondria Aspect Ratio per Image
Ilastik HT Fiji
Ilastik HT Python
Ilastik Fluo Fiji
Ilastik Fluo Python
f)
0.0216
0.9855
Mean Mitochondria Aspect Ratio per Image
Fiji Fluo
Python Fluo
Ilastik Fluo Fiji
Ilastik Fluo Python
MitoAnalyzer Fluo
MoDL Fluo
CellProfiler Fluo

**Supplementary Figure S7.** Comparison of mean mitochondrial morphological features across image analysis pipelines. (a-b) Mean mitochondrial area per image. (a) Comparison of mean mitochondrial area obtained using ilastik-based segmentation across imaging modalities (holotomographic, HT, and fluorescence, Fluo) and downstream quantification environments. (b) Comparison of mean mitochondrial area per image quantified from fluorescence images using different analysis pipelines. Only p-values > 0.01 are shown in the graph; complete statistical comparisons are reported in Supplementary Table 2. (c-d) Mean mitochondrial perimeter per image. (c) Comparison of mean mitochondrial perimeter obtained using ilastik-based segmentation across imaging modalities (holotomographic and fluorescence) and downstream quantification environments. (d) Comparison of mean mitochondrial perimeter per image quantified from fluorescence images using different analysis pipelines. Only p-values > 0.01 are shown in the graph; complete statistical comparisons are reported in Supplementary Table 3. (e-f) Mean mitochondrial aspect ratio per image. (e) Comparison of mean mitochondrial aspect ratio obtained using ilastik-based segmentation across imaging modalities (holotomographic and fluorescence) and downstream quantification environments. (f) Comparison of mean mitochondrial aspect ratio per image quantified from fluorescence images using different analysis pipelines. Only p-values > 0.01 are shown in the graph; complete statistical comparisons are reported in Supplementary Table 4. Masks derived from holotomographic segmentations using threshold-based approaches were excluded from quantitative analysis due to insufficient segmentation quality.

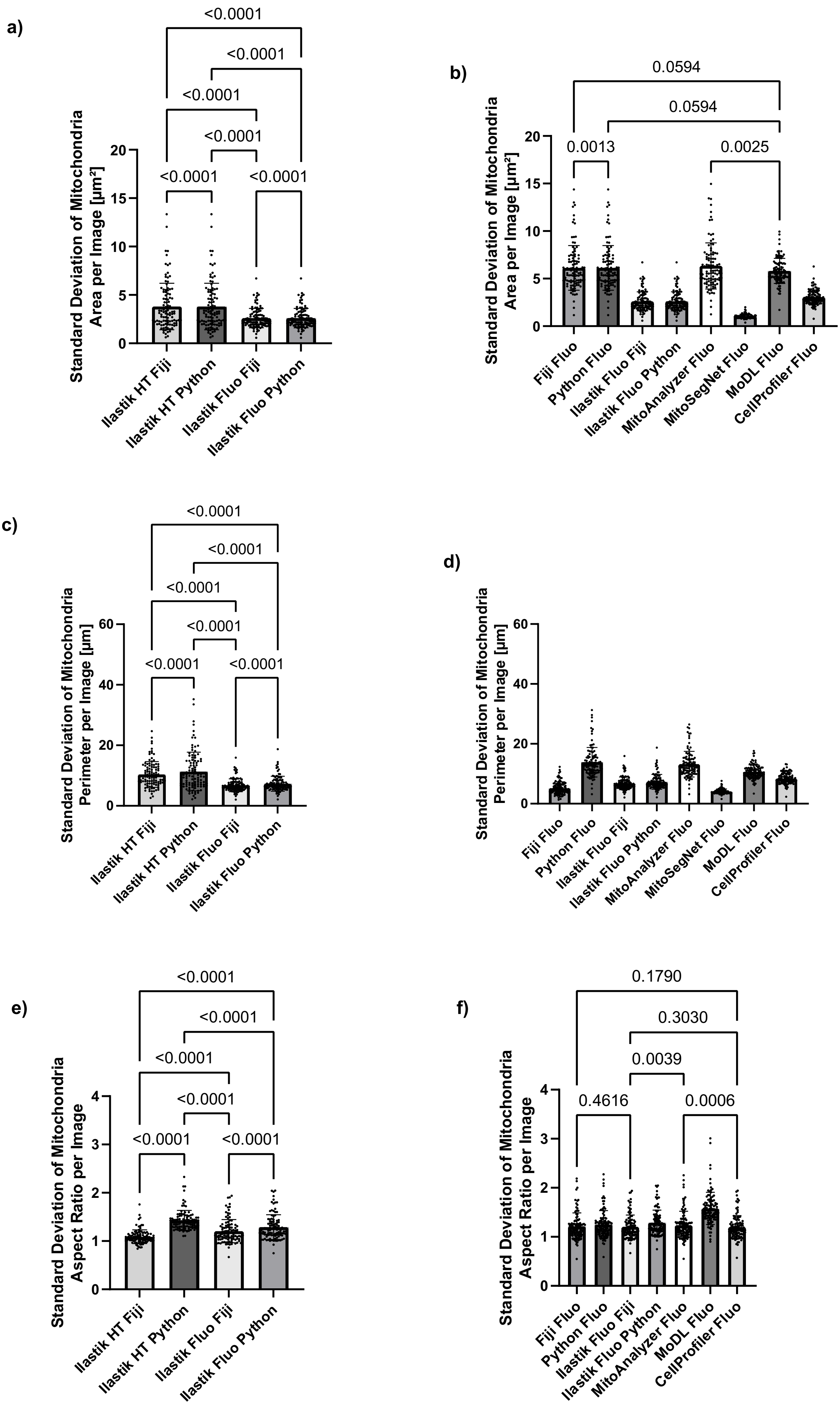
a)
<0.0001
<0.0001
<0.0001
<0.0001
<0.0001
<0.0001
Standard Deviation of Mitochondria Area per Image [µm²]
Ilastik HT Fiji
Ilastik HT Python
Ilastik Fluo Fiji
Ilastik Fluo Python
b)
0.0594
0.0594
0.0013
0.0025
Standard Deviation of Mitochondria Area per Image [µm²]
Fiji Fluo
Python Fluo
Ilastik Fluo Fiji
Ilastik Fluo Python
MitoAnalyzer Fluo
MitoSegNet Fluo
MoDL Fluo
CellProfiler Fluo
c)
<0.0001
<0.0001
<0.0001
<0.0001
<0.0001
<0.0001
Standard Deviation of Mitochondria Perimeter per Image [µm]
Ilastik HT Fiji
Ilastik HT Python
Ilastik Fluo Fiji
Ilastik Fluo Python
d)
Standard Deviation of Mitochondria Perimeter per Image [µm]
Fiji Fluo
Python Fluo
Ilastik Fluo Fiji
Ilastik Fluo Python
MitoAnalyzer Fluo
MitoSegNet Fluo
MoDL Fluo
CellProfiler Fluo
e)
<0.0001
<0.0001
<0.0001
<0.0001
<0.0001
<0.0001
Standard Deviation of Mitochondria Aspect Ratio per Image
Ilastik HT Fiji
Ilastik HT Python
Ilastik Fluo Fiji
Ilastik Fluo Python
f)
0.1790
0.3030
0.0039
0.4616
0.0006
Standard Deviation of Mitochondria Aspect Ratio per Image
Fiji Fluo
Python Fluo
Ilastik Fluo Fiji
Ilastik Fluo Python
MitoAnalyzer Fluo
MoDL Fluo
CellProfiler Fluo

**Supplementary Figure S8.** Comparison of mitochondrial feature variability across image analysis pipelines. (a-b) Within-image variability of mitochondrial area, quantified as the standard deviation per image. (a) Comparison of mitochondrial area variability obtained using ilastik-based segmentation across imaging modalities (holotomographic, HT, and fluorescence, Fluo) and downstream quantification environments. (b) Comparison of mitochondrial area variability per image quantified from fluorescence images using different analysis pipelines. Only p-values > 0.01 are shown in the graph; complete statistical comparisons are reported in Supplementary Table 5. (c-d) Within-image variability of mitochondrial perimeter, quantified as the standard deviation per image. (c) Comparison of mitochondrial perimeter variability obtained using ilastik-based segmentation across imaging modalities (holotomographic and fluorescence) and downstream quantification environments. (d) Comparison of mitochondrial perimeter variability per image quantified from fluorescence images using different analysis pipelines. Only p-values > 0.01 are shown in the graph; complete statistical comparisons are reported in Supplementary Table 6. (e-f) Within-image variability of mitochondrial aspect ratio, quantified as the standard deviation per image. (e) Comparison of mitochondrial aspect ratio variability obtained using ilastik-based segmentation across imaging modalities (holotomographic and fluorescence) and downstream quantification environments. (f) Comparison of mitochondrial aspect ratio variability per image quantified from fluorescence images using different analysis pipelines. Only p-values > 0.01 are shown in the graph; complete statistical comparisons are reported in Supplementary Table 7. Masks derived from holotomographic segmentations using threshold-based approaches were excluded from quantitative analysis due to insufficient segmentation quality.

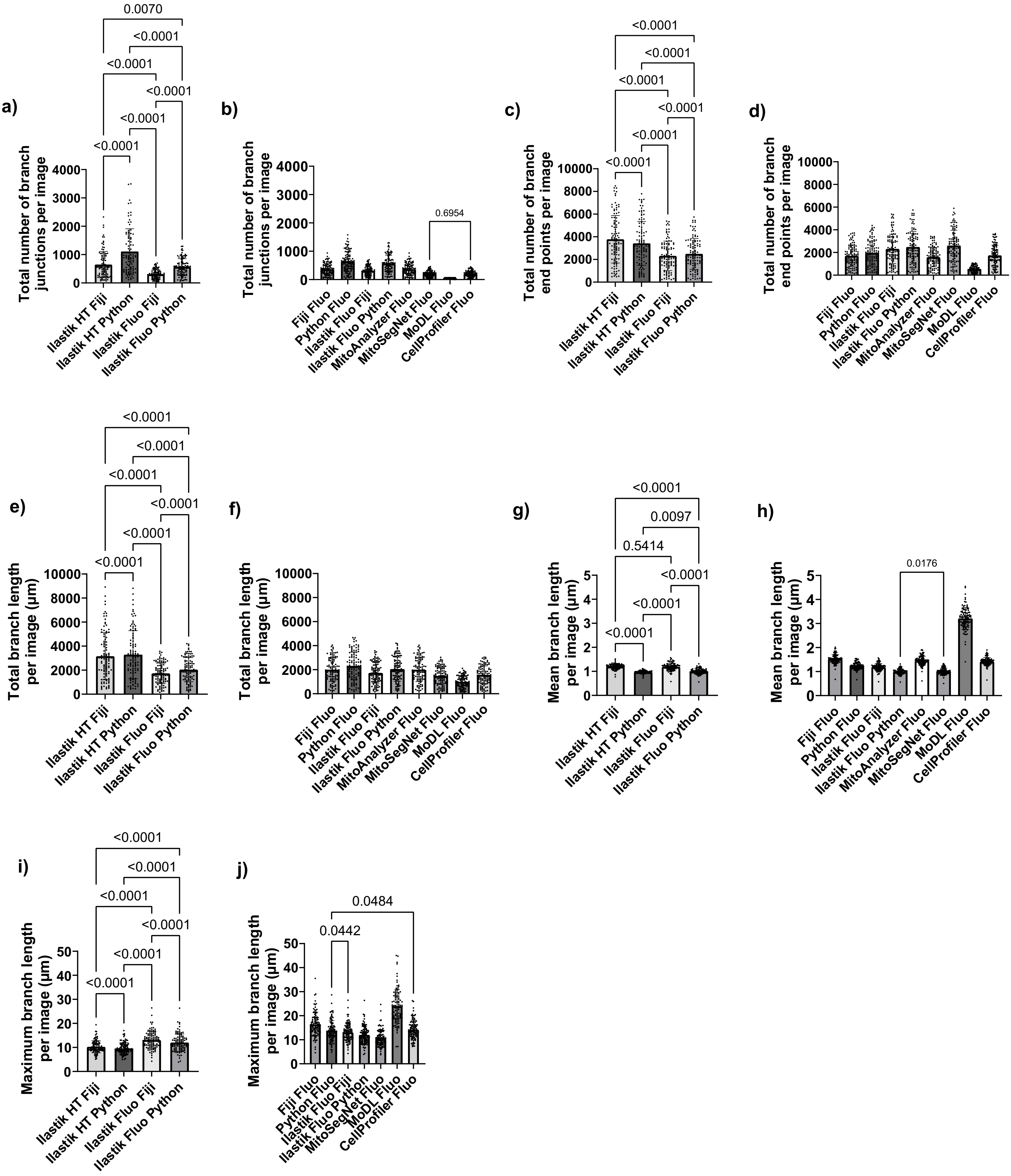
a)
Total number of branch junctions per image
Ilastik HT Fiji
Ilastik HT Python
Ilastik Fluo Fiji
Ilastik Fluo Python
0.0070
<0.0001
b)
Total number of branch junctions per image
Fiji Fluo
Python Fluo
Ilastik Fluo Fiji
Ilastik Fluo Python
MitoAnalyzer Fluo
MitoSegNet Fluo
MoDL Fluo
CellProfiler Fluo
0.6954
c)
Total number of branch end points per image
d)
Total number of branch end points per image
e)
Total branch length per image (µm)
f)
Total branch length per image (µm)
g)
Mean branch length per image (µm)
0.0097
0.5414
h)
Mean branch length per image (µm)
0.0176
i)
Maximum branch length per image (µm)
j)
Maximum branch length per image (µm)
0.0484
0.0442

**Supplementary Figure S9**. Comparison of mitochondrial network quantitative analysis across image analysis pipelines. (a-b) Total number of mitochondrial branch junctions per image. (a) Comparison of branch junction counts obtained using ilastik-based segmentation across imaging modalities (holotomographic, HT, and fluorescence, Fluo) and downstream skeletonization environments. (b) Comparison of branch junction counts per image quantified from fluorescence images using different analysis pipelines. Statistical comparisons are reported in Supplementary Table 9. (c-d) Total number of mitochondrial branch end points per image. (c) Comparison of branch end point counts obtained using ilastik-based segmentation across imaging modalities (holotomographic and fluorescence) and downstream skeletonization environments. (d) Comparison of branch end point counts per image quantified from fluorescence images using different analysis pipelines. Statistical comparisons are reported in Supplementary Table 10. (e-f) Total mitochondrial branch length per image. (e) Comparison of total branch length obtained using ilastik-based segmentation across imaging modalities (holotomographic and fluorescence) and downstream skeletonization environments. (f) Comparison of total branch length per image quantified from fluorescence images using different analysis pipelines. Statistical comparisons are reported in Supplementary Table 11. (g-h) Mean mitochondrial branch length per image. (g) Comparison of mean branch length obtained using ilastik-based segmentation across imaging modalities (holotomographic and fluorescence) and downstream skeletonization environments. (h) Comparison of mean branch length per image quantified from fluorescence images using different analysis pipelines. Statistical comparisons are reported in Supplementary Table 12. (i-j) Maximum mitochondrial branch length per image. (i) Comparison of maximum branch length obtained using ilastik-based segmentation across imaging modalities (holotomographic and fluorescence) and downstream skeletonization environments. (j) Comparison of maximum branch length per image quantified from fluorescence images using different analysis pipelines. Statistical comparisons are reported in

Supplementary Table 13. Skeletons derived from holotomographic segmentations using threshold-based approaches were excluded from quantitative analysis due to insufficient segmentation quality for reliable network quantification.

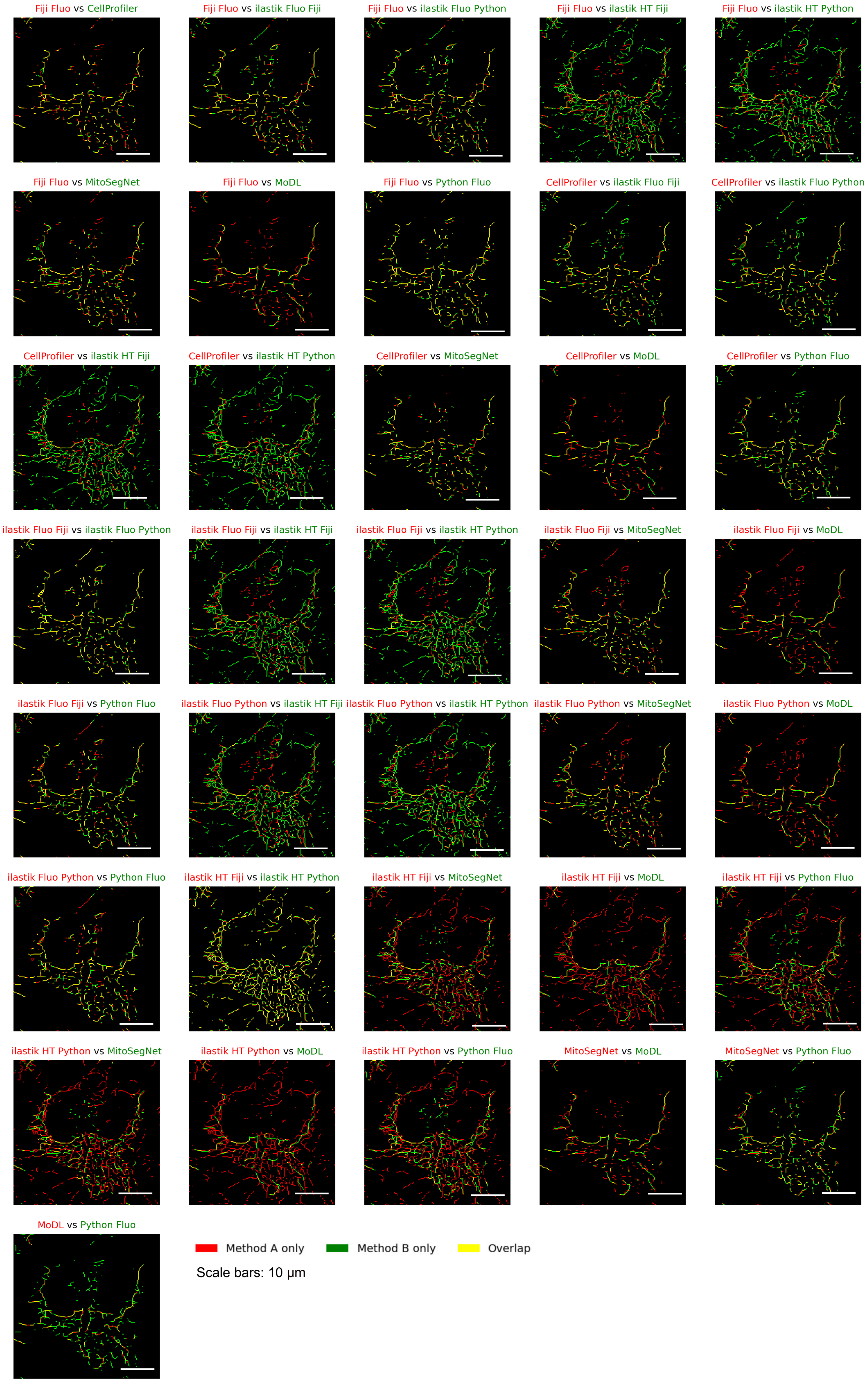

Fiji Fluo vs CellProfiler
Fiji Fluo vs ilastik Fluo Fiji
Fiji Fluo vs ilastik Fluo Python
Fiji Fluo vs ilastik HT Fiji
Fiji Fluo vs ilastik HT Python
Fiji Fluo vs MitoSegNet
Fiji Fluo vs MoDL
Fiji Fluo vs Python Fluo
CellProfiler vs ilastik Fluo Fiji
CellProfiler vs ilastik Fluo Python
CellProfiler vs ilastik HT Fiji
CellProfiler vs ilastik HT Python
CellProfiler vs MitoSegNet
CellProfiler vs MoDL
CellProfiler vs Python Fluo
ilastik Fluo Fiji vs ilastik Fluo Python
ilastik Fluo Fiji vs ilastik HT Fiji
ilastik Fluo Fiji vs ilastik HT Python
ilastik Fluo Fiji vs MitoSegNet
ilastik Fluo Fiji vs MoDL
ilastik Fluo Fiji vs Python Fluo
ilastik Fluo Python vs ilastik HT Fiji
ilastik Fluo Python vs ilastik HT Python
ilastik Fluo Python vs MitoSegNet
ilastik Fluo Python vs MoDL
ilastik Fluo Python vs Python Fluo
ilastik HT Fiji vs ilastik HT Python
ilastik HT Fiji vs MitoSegNet
ilastik HT Fiji vs MoDL
ilastik HT Fiji vs Python Fluo
ilastik HT Python vs MitoSegNet
ilastik HT Python vs MoDL
ilastik HT Python vs Python Fluo
MitoSegNet vs MoDL
MitoSegNet vs Python Fluo
MoDL vs Python Fluo
Method A only
Method B only
Overlap
Scale bars: 10 μm

**Supplementary Figure S10.** Visual comparison of mitochondrial skeleton overlaps across analysis pipelines. Pairwise overlays of mitochondrial skeletons generated by different analysis pipelines applied to the same fluorescence image (reproduced from Figure 5 for consistency). For each panel, skeletons from two pipelines are overlaid to visually illustrate spatial agreement and disagreement between methods. Skeleton pixels detected only by the first pipeline are shown in red, pixels detected only by the second pipeline are shown in green, and pixels shared by both pipelines (overlap) are shown in yellow. High proportions of yellow indicate strong spatial agreement, whereas increased red or green signal reflects method-specific detections or fragmentation. This qualitative comparison was used as an initial assessment of agreement between pipelines and motivated the subsequent quantitative evaluation of skeleton overlap using Dice and relaxed Dice coefficients (Figure 7).

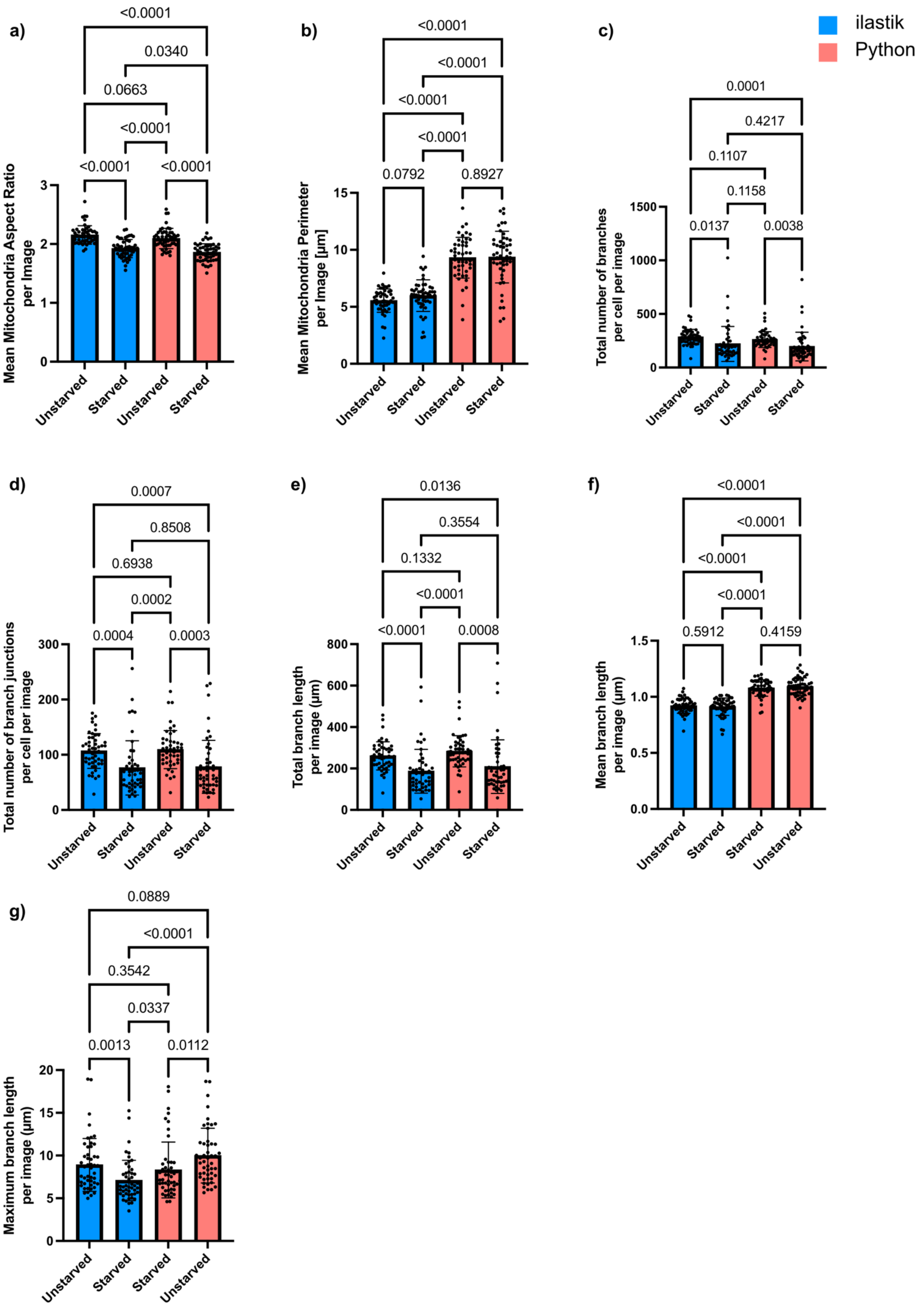
ilastik
Python
a)
<0.0001
0.0340
0.0663
<0.0001
<0.0001
<0.0001
Mean Mitochondria Aspect Ratio per Image
Unstarved
Starved
Unstarved
Starved
b)
<0.0001
<0.0001
<0.0001
<0.0001
0.0792
0.8927
Mean Mitochondria Perimeter per Image [µm]
Unstarved
Starved
Unstarved
Starved
c)
0.0001
0.4217
0.1107
0.1158
0.0137
0.0038
Total number of branches per cell per image
Unstarved
Starved
Unstarved
Starved
d)
0.0007
0.8508
0.6938
0.0002
0.0004
0.0003
Total number of branch junctions per cell per image
Unstarved
Starved
Unstarved
Starved
e)
0.0136
0.3554
0.1332
<0.0001
<0.0001
0.0008
Total branch length per image (µm)
Unstarved
Starved
Unstarved
Starved
f)
<0.0001
<0.0001
<0.0001
<0.0001
0.5912
0.4159
Mean branch length per image (µm)
Unstarved
Starved
Starved
Unstarved
g)
0.0889
<0.0001
0.3542
0.0337
0.0013
0.0112
Maximum branch length per image (µm)
Unstarved
Starved
Starved
Unstarved

Supplementary Figure S11: Comparison of ilastik- and Python-based quantification of mitochondrial morphology and network architecture. The mitochondrial features were either analyzed with ilastik (blue) or Python (red). (a) Mean mitochondrial aspect ratio, (b) mean mitochondrial perimeter, (c) total number of branches, (d) total number of branch junctions, (e) total branch length, (f) mean branch length, (g) maximum branch length. Statistical analyses: Welch's ANOVA.

**Supplementary Table 1:** Number of Mitochodria per image (Figure 4c)

| Comparison | P-value |
|---|---|
| Fiji Fluo vs. Python Fluo | <0.0001 |
| Fiji Fluo vs. Ilastik Fluo Fiji | <0.0001 |
| Fiji Fluo vs. Ilastik Fluo Python | <0.0001 |
| Fiji Fluo vs. MitoAnalyzer Fluo | <0.0001 |
| Fiji Fluo vs. MitoSegNet Fluo | <0.0001 |
| Fiji Fluo vs. MoDL Fluo | <0.0001 |
| Fiji Fluo vs. CellProfiler Fluo | <0.0001 |
| Python Fluo vs. Ilastik Fluo Fiji | <0.0001 |
| Python Fluo vs. Ilastik Fluo Python | <0.0001 |
| Python Fluo vs. MitoAnalyzer Fluo | <0.0001 |
| Python Fluo vs. MitoSegNet Fluo | <0.0001 |
| Python Fluo vs. MoDL Fluo | <0.0001 |
| Python Fluo vs. CellProfiler Fluo | 0.3197 |
| Ilastik Fluo Fiji vs. Ilastik Fluo Python | <0.0001 |
| Ilastik Fluo Fiji vs. MitoAnalyzer Fluo | <0.0001 |
| Ilastik Fluo Fiji vs. MitoSegNet Fluo | <0.0001 |
| Ilastik Fluo Fiji vs. MoDL Fluo | <0.0001 |
| Ilastik Fluo Fiji vs. CellProfiler Fluo | <0.0001 |
| Ilastik Fluo Python vs. MitoAnalyzer Fluo | <0.0001 |
| Ilastik Fluo Python vs. MitoSegNet Fluo | <0.0001 |
| Ilastik Fluo Python vs. MoDL Fluo | <0.0001 |
| Ilastik Fluo Python vs. CellProfiler Fluo | <0.0001 |
| MitoAnalyzer Fluo vs. MitoSegNet Fluo | <0.0001 |
| MitoAnalyzer Fluo vs. MoDL Fluo | <0.0001 |
| MitoAnalyzer Fluo vs. CellProfiler Fluo | <0.0001 |
| MitoSegNet Fluo vs. MoDL Fluo | <0.0001 |
| MitoSegNet Fluo vs. CellProfiler Fluo | <0.0001 |
| MoDL Fluo vs. CellProfiler Fluo | <0.0001 |

**Supplementary Table 2:** Mean Mitochodria Area per image (Supp. Figure 7b)

| Comparison | P-value |
|---|---|
| Fiji Fluo vs. Python Fluo | <0.0001 |
| Fiji Fluo vs. Ilastik Fluo Fiji | <0.0001 |
| Fiji Fluo vs. Ilastik Fluo Python | <0.0001 |
| Fiji Fluo vs. MitoAnalyzer Fluo | <0.0001 |
| Fiji Fluo vs. MitoSegNet Fluo | <0.0001 |
| Fiji Fluo vs. MoDL Fluo | <0.0001 |
| Fiji Fluo vs. CellProfiler Fluo | <0.0001 |
| Python Fluo vs. Ilastik Fluo Fiji | <0.0001 |
| Python Fluo vs. Ilastik Fluo Python | <0.0001 |
| Python Fluo vs. MitoAnalyzer Fluo | <0.0001 |
| Python Fluo vs. MitoSegNet Fluo | <0.0001 |
| Python Fluo vs. MoDL Fluo | <0.0001 |
| Python Fluo vs. CellProfiler Fluo | <0.0001 |
| Ilastik Fluo Fiji vs. Ilastik Fluo Python | <0.0001 |
| Ilastik Fluo Fiji vs. MitoAnalyzer Fluo | <0.0001 |
| Ilastik Fluo Fiji vs. MitoSegNet Fluo | <0.0001 |
| Ilastik Fluo Fiji vs. MoDL Fluo | <0.0001 |
| Ilastik Fluo Fiji vs. CellProfiler Fluo | <0.0001 |
| Ilastik Fluo Python vs. MitoAnalyzer Fluo | <0.0001 |
| Ilastik Fluo Python vs. MitoSegNet Fluo | <0.0001 |
| Ilastik Fluo Python vs. MoDL Fluo | <0.0001 |
| Ilastik Fluo Python vs. CellProfiler Fluo | <0.0001 |
| MitoAnalyzer Fluo vs. MitoSegNet Fluo | <0.0001 |
| MitoAnalyzer Fluo vs. MoDL Fluo | <0.0001 |
| MitoAnalyzer Fluo vs. CellProfiler Fluo | <0.0001 |
| MitoSegNet Fluo vs. MoDL Fluo | <0.0001 |
| MitoSegNet Fluo vs. CellProfiler Fluo | <0.0001 |
| MoDL Fluo vs. CellProfiler Fluo | <0.0001 |

**Supplementary Table 3:** Mean Mitochodria Perimeter per image (Supp. Figure 7d)

| Comparison | P-value |
|---|---|
| Fiji Fluo vs. Python Fluo | 0.5218 |
| Fiji Fluo vs. Ilastik Fluo Fiji | <0.0001 |
| Fiji Fluo vs. Ilastik Fluo Python | <0.0001 |
| Fiji Fluo vs. MitoAnalyzer Fluo | <0.0001 |
| Fiji Fluo vs. MitoSegNet Fluo | <0.0001 |
| Fiji Fluo vs. MoDL Fluo | <0.0001 |
| Fiji Fluo vs. CellProfiler Fluo | <0.0001 |
| Python Fluo vs. Ilastik Fluo Fiji | <0.0001 |
| Python Fluo vs. Ilastik Fluo Python | <0.0001 |
| Python Fluo vs. MitoAnalyzer Fluo | <0.0001 |
| Python Fluo vs. MitoSegNet Fluo | <0.0001 |
| Python Fluo vs. MoDL Fluo | <0.0001 |
| Python Fluo vs. CellProfiler Fluo | <0.0001 |
| Ilastik Fluo Fiji vs. Ilastik Fluo Python | <0.0001 |
| Ilastik Fluo Fiji vs. MitoAnalyzer Fluo | <0.0001 |
| Ilastik Fluo Fiji vs. MitoSegNet Fluo | <0.0001 |
| Ilastik Fluo Fiji vs. MoDL Fluo | <0.0001 |
| Ilastik Fluo Fiji vs. CellProfiler Fluo | <0.0001 |
| Ilastik Fluo Python vs. MitoAnalyzer Fluo | <0.0001 |
| Ilastik Fluo Python vs. MitoSegNet Fluo | <0.0001 |
| Ilastik Fluo Python vs. MoDL Fluo | <0.0001 |
| Ilastik Fluo Python vs. CellProfiler Fluo | <0.0001 |
| MitoAnalyzer Fluo vs. MitoSegNet Fluo | <0.0001 |
| MitoAnalyzer Fluo vs. MoDL Fluo | <0.0001 |
| MitoAnalyzer Fluo vs. CellProfiler Fluo | <0.0001 |
| MitoSegNet Fluo vs. MoDL Fluo | <0.0001 |
| MitoSegNet Fluo vs. CellProfiler Fluo | <0.0001 |
| MoDL Fluo vs. CellProfiler Fluo | <0.0001 |

**Supplementary Table 4: Mean Mitochondria Aspect Ratio per image (Supp. Figure 7f)**

| Comparison | P-value |
|---|---|
| Fiji Fluo vs. Python Fluo | <0.0001 |
| Fiji Fluo vs. Ilastik Fluo Fiji | <0.0001 |
| Fiji Fluo vs. Ilastik Fluo Python | <0.0001 |
| Fiji Fluo vs. MitoAnalyzer Fluo | <0.0001 |
| Fiji Fluo vs. MoDL Fluo | <0.0001 |
| Fiji Fluo vs. CellProfiler Fluo | <0.0001 |
| Python Fluo vs. Ilastik Fluo Fiji | <0.0001 |
| Python Fluo vs. Ilastik Fluo Python | <0.0001 |
| Python Fluo vs. MitoAnalyzer Fluo | <0.0001 |
| Python Fluo vs. MoDL Fluo | <0.0001 |
| Python Fluo vs. CellProfiler Fluo | 0.0216 |
| Ilastik Fluo Fiji vs. Ilastik Fluo Python | <0.0001 |
| Ilastik Fluo Fiji vs. MitoAnalyzer Fluo | <0.0001 |
| Ilastik Fluo Fiji vs. MoDL Fluo | <0.0001 |
| Ilastik Fluo Fiji vs. CellProfiler Fluo | <0.0001 |
| Ilastik Fluo Python vs. MitoAnalyzer Fluo | <0.0001 |
| Ilastik Fluo Python vs. MoDL Fluo | <0.0001 |
| Ilastik Fluo Python vs. CellProfiler Fluo | <0.0001 |
| MitoAnalyzer Fluo vs. MoDL Fluo | <0.0001 |
| MitoAnalyzer Fluo vs. CellProfiler Fluo | 0.9855 |
| MoDL Fluo vs. CellProfiler Fluo | <0.0001 |

**Supplementary Table 5:** Standard Deviation of Mitochodria Area per image (Supp. Figure 8b)

| Comparison | P-value |
|---|---|
| Fiji Fluo vs. Python Fluo | 0.0013 |
| Fiji Fluo vs. Ilastik Fluo Fiji | <0.0001 |
| Fiji Fluo vs. Ilastik Fluo Python | <0.0001 |
| Fiji Fluo vs. MitoAnalyzer Fluo | <0.0001 |
| Fiji Fluo vs. MitoSegNet Fluo | <0.0001 |
| Fiji Fluo vs. MoDL Fluo | 0.0594 |
| Fiji Fluo vs. CellProfiler Fluo | <0.0001 |
| Python Fluo vs. Ilastik Fluo Fiji | <0.0001 |
| Python Fluo vs. Ilastik Fluo Python | <0.0001 |
| Python Fluo vs. MitoAnalyzer Fluo | <0.0001 |
| Python Fluo vs. MitoSegNet Fluo | <0.0001 |
| Python Fluo vs. MoDL Fluo | 0.0594 |
| Python Fluo vs. CellProfiler Fluo | <0.0001 |
| Ilastik Fluo Fiji vs. Ilastik Fluo Python | <0.0001 |
| Ilastik Fluo Fiji vs. MitoAnalyzer Fluo | <0.0001 |
| Ilastik Fluo Fiji vs. MitoSegNet Fluo | <0.0001 |
| Ilastik Fluo Fiji vs. MoDL Fluo | <0.0001 |
| Ilastik Fluo Fiji vs. CellProfiler Fluo | <0.0001 |
| Ilastik Fluo Python vs. MitoAnalyzer Fluo | <0.0001 |
| Ilastik Fluo Python vs. MitoSegNet Fluo | <0.0001 |
| Ilastik Fluo Python vs. MoDL Fluo | <0.0001 |
| Ilastik Fluo Python vs. CellProfiler Fluo | <0.0001 |
| MitoAnalyzer Fluo vs. MitoSegNet Fluo | <0.0001 |
| MitoAnalyzer Fluo vs. MoDL Fluo | 0.0025 |
| MitoAnalyzer Fluo vs. CellProfiler Fluo | <0.0001 |
| MitoSegNet Fluo vs. MoDL Fluo | <0.0001 |
| MitoSegNet Fluo vs. CellProfiler Fluo | <0.0001 |
| MoDL Fluo vs. CellProfiler Fluo | <0.0001 |

**Supplementary Table 6:** Standard Deviation of Mitochodria Perimeter per image (Supp. Figure 8d)

| Comparison | P-value |
|---|---|
| Fiji Fluo vs. Python Fluo | <0.0001 |
| Fiji Fluo vs. Ilastik Fluo Fiji | <0.0001 |
| Fiji Fluo vs. Ilastik Fluo Python | <0.0001 |
| Fiji Fluo vs. MitoAnalyzer Fluo | <0.0001 |
| Fiji Fluo vs. MitoSegNet Fluo | <0.0001 |
| Fiji Fluo vs. MoDL Fluo | <0.0001 |
| Fiji Fluo vs. CellProfiler Fluo | <0.0001 |
| Python Fluo vs. Ilastik Fluo Fiji | <0.0001 |
| Python Fluo vs. Ilastik Fluo Python | <0.0001 |
| Python Fluo vs. MitoAnalyzer Fluo | <0.0001 |
| Python Fluo vs. MitoSegNet Fluo | <0.0001 |
| Python Fluo vs. MoDL Fluo | <0.0001 |
| Python Fluo vs. CellProfiler Fluo | <0.0001 |
| Ilastik Fluo Fiji vs. Ilastik Fluo Python | <0.0001 |
| Ilastik Fluo Fiji vs. MitoAnalyzer Fluo | <0.0001 |
| Ilastik Fluo Fiji vs. MitoSegNet Fluo | <0.0001 |
| Ilastik Fluo Fiji vs. MoDL Fluo | <0.0001 |
| Ilastik Fluo Fiji vs. CellProfiler Fluo | <0.0001 |
| Ilastik Fluo Python vs. MitoAnalyzer Fluo | <0.0001 |
| Ilastik Fluo Python vs. MitoSegNet Fluo | <0.0001 |
| Ilastik Fluo Python vs. MoDL Fluo | <0.0001 |
| Ilastik Fluo Python vs. CellProfiler Fluo | <0.0001 |
| MitoAnalyzer Fluo vs. MitoSegNet Fluo | <0.0001 |
| MitoAnalyzer Fluo vs. MoDL Fluo | <0.0001 |
| MitoAnalyzer Fluo vs. CellProfiler Fluo | <0.0001 |
| MitoSegNet Fluo vs. MoDL Fluo | <0.0001 |
| MitoSegNet Fluo vs. CellProfiler Fluo | <0.0001 |
| MoDL Fluo vs. CellProfiler Fluo | <0.0001 |

**Supplementary Table 7:** Standard Deviation of Mitochodria Aspect Ratio per image (Supp. Figure 8f)

| Comparison | P-value |
|---|---|
| Fiji Fluo vs. Python Fluo | <0.0001 |
| Fiji Fluo vs. Ilastik Fluo Fiji | 0.4616 |
| Fiji Fluo vs. Ilastik Fluo Python | <0.0001 |
| Fiji Fluo vs. MitoAnalyzer Fluo | <0.0001 |
| Fiji Fluo vs. MoDL Fluo | <0.0001 |
| Fiji Fluo vs. CellProfiler Fluo | 0.179 |
| Python Fluo vs. Ilastik Fluo Fiji | <0.0001 |
| Python Fluo vs. Ilastik Fluo Python | 0.0015 |
| Python Fluo vs. MitoAnalyzer Fluo | <0.0001 |
| Python Fluo vs. MoDL Fluo | <0.0001 |
| Python Fluo vs. CellProfiler Fluo | <0.0001 |
| Ilastik Fluo Fiji vs. Ilastik Fluo Python | <0.0001 |
| Ilastik Fluo Fiji vs. MitoAnalyzer Fluo | 0.0039 |
| Ilastik Fluo Fiji vs. MoDL Fluo | <0.0001 |
| Ilastik Fluo Fiji vs. CellProfiler Fluo | 0.303 |
| Ilastik Fluo Python vs. MitoAnalyzer Fluo | <0.0001 |
| Ilastik Fluo Python vs. MoDL Fluo | <0.0001 |
| Ilastik Fluo Python vs. CellProfiler Fluo | <0.0001 |
| MitoAnalyzer Fluo vs. MoDL Fluo | <0.0001 |
| MitoAnalyzer Fluo vs. CellProfiler Fluo | 0.0006 |
| MoDL Fluo vs. CellProfiler Fluo | <0.0001 |

**Supplementary Table 8:** Total number of branches per image (Figure 6c)

| Comparison | P-value |
|---|---|
| Fiji Fluo vs. Python Fluo | <0.0001 |
| Fiji Fluo vs. Ilastik Fluo Fiji | <0.0001 |
| Fiji Fluo vs. Ilastik Fluo Python | <0.0001 |
| Fiji Fluo vs. MitoAnalyzer Fluo | <0.0001 |
| Fiji Fluo vs. MitoSegNet Fluo | <0.0001 |
| Fiji Fluo vs. MoDL Fluo | <0.0001 |
| Fiji Fluo vs. CellProfiler Fluo | <0.0001 |
| Python Fluo vs. Ilastik Fluo Fiji | <0.0001 |
| Python Fluo vs. Ilastik Fluo Python | <0.0001 |
| Python Fluo vs. MitoAnalyzer Fluo | <0.0001 |
| Python Fluo vs. MitoSegNet Fluo | <0.0001 |
| Python Fluo vs. MoDL Fluo | <0.0001 |
| Python Fluo vs. CellProfiler Fluo | <0.0001 |
| Ilastik Fluo Fiji vs. Ilastik Fluo Python | <0.0001 |
| Ilastik Fluo Fiji vs. MitoAnalyzer Fluo | <0.0001 |
| Ilastik Fluo Fiji vs. MitoSegNet Fluo | 0.5726 |
| Ilastik Fluo Fiji vs. MoDL Fluo | <0.0001 |
| Ilastik Fluo Fiji vs. CellProfiler Fluo | <0.0001 |
| Ilastik Fluo Python vs. MitoAnalyzer Fluo | <0.0001 |
| Ilastik Fluo Python vs. MitoSegNet Fluo | <0.0001 |
| Ilastik Fluo Python vs. MoDL Fluo | <0.0001 |
| Ilastik Fluo Python vs. CellProfiler Fluo | <0.0001 |
| MitoAnalyzer Fluo vs. MitoSegNet Fluo | <0.0001 |
| MitoAnalyzer Fluo vs. MoDL Fluo | <0.0001 |
| MitoAnalyzer Fluo vs. CellProfiler Fluo | <0.0001 |
| MitoSegNet Fluo vs. MoDL Fluo | <0.0001 |
| MitoSegNet Fluo vs. CellProfiler Fluo | <0.0001 |
| MoDL Fluo vs. CellProfiler Fluo | <0.0001 |

**Supplementary Table 9:** Total number of branch junctions per image (Supp. Figure 9b)

| Comparison | P-value |
|---|---|
| Fiji Fluo vs. Python Fluo | <0.0001 |
| Fiji Fluo vs. Ilastik Fluo Fiji | <0.0001 |
| Fiji Fluo vs. Ilastik Fluo Python | <0.0001 |
| Fiji Fluo vs. MitoAnalyzer Fluo | No diff. |
| Fiji Fluo vs. MitoSegNet Fluo | <0.0001 |
| Fiji Fluo vs. MoDL Fluo | <0.0001 |
| Fiji Fluo vs. CellProfiler Fluo | <0.0001 |
| Python Fluo vs. Ilastik Fluo Fiji | <0.0001 |
| Python Fluo vs. Ilastik Fluo Python | <0.0001 |
| Python Fluo vs. MitoAnalyzer Fluo | <0.0001 |
| Python Fluo vs. MitoSegNet Fluo | <0.0001 |
| Python Fluo vs. MoDL Fluo | <0.0001 |
| Python Fluo vs. CellProfiler Fluo | <0.0001 |
| Ilastik Fluo Fiji vs. Ilastik Fluo Python | <0.0001 |
| Ilastik Fluo Fiji vs. MitoAnalyzer Fluo | <0.0001 |
| Ilastik Fluo Fiji vs. MitoSegNet Fluo | <0.0001 |
| Ilastik Fluo Fiji vs. MoDL Fluo | <0.0001 |
| Ilastik Fluo Fiji vs. CellProfiler Fluo | <0.0001 |
| Ilastik Fluo Python vs. MitoAnalyzer Fluo | <0.0001 |
| Ilastik Fluo Python vs. MitoSegNet Fluo | <0.0001 |
| Ilastik Fluo Python vs. MoDL Fluo | <0.0001 |
| Ilastik Fluo Python vs. CellProfiler Fluo | <0.0001 |
| MitoAnalyzer Fluo vs. MitoSegNet Fluo | <0.0001 |
| MitoAnalyzer Fluo vs. MoDL Fluo | <0.0001 |
| MitoAnalyzer Fluo vs. CellProfiler Fluo | <0.0001 |
| MitoSegNet Fluo vs. MoDL Fluo | <0.0001 |
| MitoSegNet Fluo vs. CellProfiler Fluo | 0.6954 |
| MoDL Fluo vs. CellProfiler Fluo | <0.0001 |

**Supplementary Table 10:** Total number of branch end points per image (Supp. Figure 9d)

| Comparison | P-value |
|---|---|
| Fiji Fluo vs. Python Fluo | <0,0001 |
| Fiji Fluo vs. Ilastik Fluo Fiji | <0,0001 |
| Fiji Fluo vs. Ilastik Fluo Python | <0,0001 |
| Fiji Fluo vs. MitoAnalyzer Fluo | <0,0001 |
| Fiji Fluo vs. MitoSegNet Fluo | <0,0001 |
| Fiji Fluo vs. MoDL Fluo | <0,0001 |
| Fiji Fluo vs. CellProfiler Fluo | <0,0001 |
| Python Fluo vs. Ilastik Fluo Fiji | <0,0001 |
| Python Fluo vs. Ilastik Fluo Python | <0,0001 |
| Python Fluo vs. MitoAnalyzer Fluo | <0,0001 |
| Python Fluo vs. MitoSegNet Fluo | <0,0001 |
| Python Fluo vs. MoDL Fluo | <0,0001 |
| Python Fluo vs. CellProfiler Fluo | <0,0001 |
| Ilastik Fluo Fiji vs. Ilastik Fluo Python | <0,0001 |
| Ilastik Fluo Fiji vs. MitoAnalyzer Fluo | <0,0001 |
| Ilastik Fluo Fiji vs. MitoSegNet Fluo | <0,0001 |
| Ilastik Fluo Fiji vs. MoDL Fluo | <0,0001 |
| Ilastik Fluo Fiji vs. CellProfiler Fluo | <0,0001 |
| Ilastik Fluo Python vs. MitoAnalyzer Fluo | <0,0001 |
| Ilastik Fluo Python vs. MitoSegNet Fluo | <0,0001 |
| Ilastik Fluo Python vs. MoDL Fluo | <0,0001 |
| Ilastik Fluo Python vs. CellProfiler Fluo | <0,0001 |
| MitoAnalyzer Fluo vs. MitoSegNet Fluo | <0,0001 |
| MitoAnalyzer Fluo vs. MoDL Fluo | <0,0001 |
| MitoAnalyzer Fluo vs. CellProfiler Fluo | <0,0001 |
| MitoSegNet Fluo vs. MoDL Fluo | <0,0001 |
| MitoSegNet Fluo vs. CellProfiler Fluo | <0,0001 |
| MoDL Fluo vs. CellProfiler Fluo | <0,0001 |

**Supplementary Table 11:** Total branch length per image (Supp. Figure 9f)

| Comparison | P-value |
|---|---|
| Fiji Fluo vs. Python Fluo | <0.0001 |
| Fiji Fluo vs. Ilastik Fluo Fiji | <0.0001 |
| Fiji Fluo vs. Ilastik Fluo Python | <0.0001 |
| Fiji Fluo vs. MitoAnalyzer Fluo | <0.0001 |
| Fiji Fluo vs. MitoSegNet Fluo | <0.0001 |
| Fiji Fluo vs. MoDL Fluo | <0.0001 |
| Fiji Fluo vs. CellProfiler Fluo | 0.001 |
| Python Fluo vs. Ilastik Fluo Fiji | <0.0001 |
| Python Fluo vs. Ilastik Fluo Python | <0.0001 |
| Python Fluo vs. MitoAnalyzer Fluo | <0.0001 |
| Python Fluo vs. MitoSegNet Fluo | <0.0001 |
| Python Fluo vs. MoDL Fluo | <0.0001 |
| Python Fluo vs. CellProfiler Fluo | <0.0001 |
| Ilastik Fluo Fiji vs. Ilastik Fluo Python | <0.0001 |
| Ilastik Fluo Fiji vs. MitoAnalyzer Fluo | <0.0001 |
| Ilastik Fluo Fiji vs. MitoSegNet Fluo | <0.0001 |
| Ilastik Fluo Fiji vs. MoDL Fluo | <0.0001 |
| Ilastik Fluo Fiji vs. CellProfiler Fluo | <0.0001 |
| Ilastik Fluo Python vs. MitoAnalyzer Fluo | <0.0001 |
| Ilastik Fluo Python vs. MitoSegNet Fluo | <0.0001 |
| Ilastik Fluo Python vs. MoDL Fluo | <0.0001 |
| Ilastik Fluo Python vs. CellProfiler Fluo | <0.0001 |
| MitoAnalyzer Fluo vs. MitoSegNet Fluo | <0.0001 |
| MitoAnalyzer Fluo vs. MoDL Fluo | <0.0001 |
| MitoAnalyzer Fluo vs. CellProfiler Fluo | <0.0001 |
| MitoSegNet Fluo vs. MoDL Fluo | <0.0001 |
| MitoSegNet Fluo vs. CellProfiler Fluo | <0.0001 |
| MoDL Fluo vs. CellProfiler Fluo | <0.0001 |

**Supplementary Table 12:** Mean branch length per image (μm) (Supp. Figure 9h)

| Comparison | P-value |
|---|---|
| Fiji Fluo vs. Python Fluo | <0.0001 |
| Fiji Fluo vs. Ilastik Fluo Fiji | <0.0001 |
| Fiji Fluo vs. Ilastik Fluo Python | <0.0001 |
| Fiji Fluo vs. MitoAnalyzer Fluo | <0.0001 |
| Fiji Fluo vs. MitoSegNet Fluo | <0.0001 |
| Fiji Fluo vs. MoDL Fluo | <0.0001 |
| Fiji Fluo vs. CellProfiler Fluo | <0.0001 |
| Python Fluo vs. Ilastik Fluo Fiji | <0.0001 |
| Python Fluo vs. Ilastik Fluo Python | <0.0001 |
| Python Fluo vs. MitoAnalyzer Fluo | <0.0001 |
| Python Fluo vs. MitoSegNet Fluo | <0.0001 |
| Python Fluo vs. MoDL Fluo | <0.0001 |
| Python Fluo vs. CellProfiler Fluo | <0.0001 |
| Ilastik Fluo Fiji vs. Ilastik Fluo Python | <0.0001 |
| Ilastik Fluo Fiji vs. MitoAnalyzer Fluo | <0.0001 |
| Ilastik Fluo Fiji vs. MitoSegNet Fluo | <0.0001 |
| Ilastik Fluo Fiji vs. MoDL Fluo | <0.0001 |
| Ilastik Fluo Fiji vs. CellProfiler Fluo | <0.0001 |
| Ilastik Fluo Python vs. MitoAnalyzer Fluo | <0.0001 |
| Ilastik Fluo Python vs. MitoSegNet Fluo | 0.0176 |
| Ilastik Fluo Python vs. MoDL Fluo | <0.0001 |
| Ilastik Fluo Python vs. CellProfiler Fluo | <0.0001 |
| MitoAnalyzer Fluo vs. MitoSegNet Fluo | <0.0001 |
| MitoAnalyzer Fluo vs. MoDL Fluo | <0.0001 |
| MitoAnalyzer Fluo vs. CellProfiler Fluo | <0.0001 |
| MitoSegNet Fluo vs. MoDL Fluo | <0.0001 |
| MitoSegNet Fluo vs. CellProfiler Fluo | <0.0001 |
| MoDL Fluo vs. CellProfiler Fluo | <0.0001 |

**Supplementary Table 13:** Maximum branch length per image (μm) (Supp. Figure 9j)

| Comparison | P-value |
|---|---|
| Fiji Fluo vs. Python Fluo | <0.0001 |
| Fiji Fluo vs. Ilastik Fluo Fiji | <0.0001 |
| Fiji Fluo vs. Ilastik Fluo Python | <0.0001 |
| Fiji Fluo vs. MitoSegNet Fluo | <0.0001 |
| Fiji Fluo vs. MoDL Fluo | <0.0001 |
| Fiji Fluo vs. CellProfiler Fluo | <0.0001 |
| Python Fluo vs. Ilastik Fluo Fiji | 0.0442 |
| Python Fluo vs. Ilastik Fluo Python | <0.0001 |
| Python Fluo vs. MitoSegNet Fluo | <0.0001 |
| Python Fluo vs. MoDL Fluo | <0.0001 |
| Python Fluo vs. CellProfiler Fluo | 0.0484 |
| Ilastik Fluo Fiji vs. Ilastik Fluo Python | <0.0001 |
| Ilastik Fluo Fiji vs. MitoSegNet Fluo | <0.0001 |
| Ilastik Fluo Fiji vs. MoDL Fluo | <0.0001 |
| Ilastik Fluo Fiji vs. CellProfiler Fluo | <0.0001 |
| Ilastik Fluo Python vs. MitoSegNet Fluo | <0.0001 |
| Ilastik Fluo Python vs. MoDL Fluo | <0.0001 |
| Ilastik Fluo Python vs. CellProfiler Fluo | <0.0001 |
| MitoSegNet Fluo vs. MoDL Fluo | <0.0001 |
| MitoSegNet Fluo vs. CellProfiler Fluo | <0.0001 |
| MoDL Fluo vs. CellProfiler Fluo | <0.0001 |
